# Differentiating *Bacillus subtilis* incorporates valine and methionine carbon into the backbone of specific fatty acids

**DOI:** 10.1101/2023.04.14.536918

**Authors:** Gerald E. Rowe, Jonathan Perreault

**Author notes:** **Data Availability Statement:** The data that support the findings of this study are available from the corresponding author upon reasonable request. **Funding Statement:** Funding to J. Perreault was provided by the Natural Sciences and Engineering Research Council of Canada (NSERC), Discovery Grant # 418240 and RGPIN-2019-06403. J. Perreault is a research scholar junior 2 from Fonds de Recherche du Québec en Santé (FRQS). We thank Éric Déziel for supporting early stages of this work via NSERC Discovery Grant # RGPIN-2015-03931. **Conflict of Interest Disclosure**: The authors declare no conflict of interest.

## Abstract

**Summary:** Sporulation in *Bacillus subtilis* has long been a model of cellular differentiation, many aspects of which are well understood. The early stage of this process is of particular interest, especially the interrelationship of regulatory processes with metabolism in response to environmental changes. We analyzed cellular fatty acids as their methyl esters using capillary gas chromatography coupled to mass spectrometry during the transition from vegetative growth to early sporulation phase. Measurement of changes in the content of heavy fatty acid analogs in cultures supplemented with deuterium-labeled valine or methionine, or ^13^C-labeld valine, showed that label was incorporated into the backbone of 12-methyltridecanoic and 14-methylpentadecanoic acid, in both sporulating and *Δspo0A* cultures. These fatty acids were formed starting with isobutyryl-CoA apparently originating only from *L*-valine-d_8_ in cultures so supplemented. Our observations indicate that following vegetative growth a pathway exists from certain amino acids into fatty acid methylene groups, evidently passing through propionyl-CoA. This finding has the potential to deepen understanding of the metabolic basis of cellular differentiation and identify new targets for antibiotics. We also observed a significant, continuous increase in the proportion of 13-methyltetradecanoic acid in fatty acids during the same period in which the pre-spore membrane would be formed.

*Graphical abstract:* 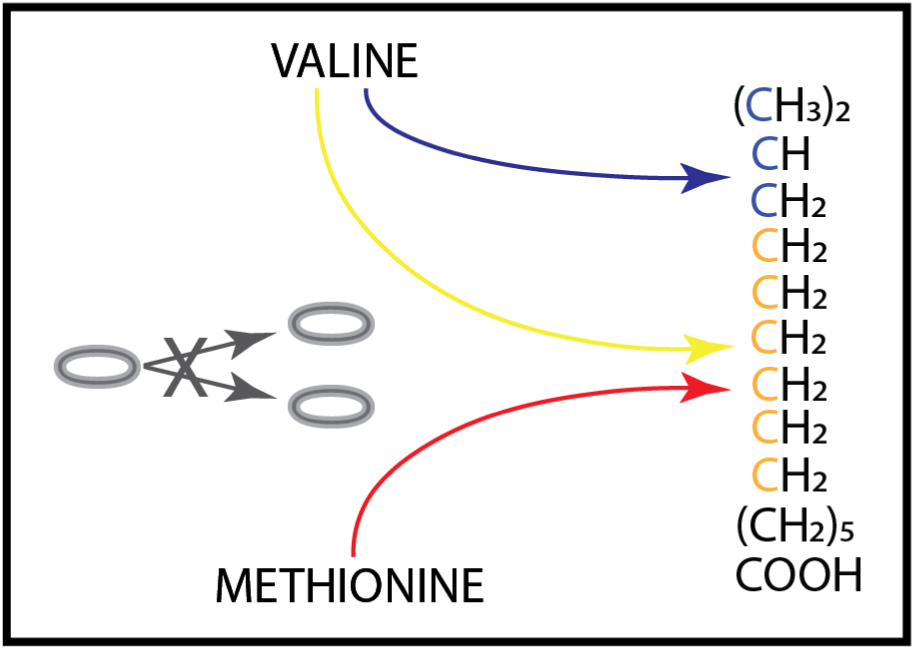

*Abbreviated Summary:* Early in at least some *Bacillus subtilis* differentiation scenarios, a straight-chain metabolite derived from propionyl-CoA is incorporated into fatty acids primed with isobutyryl-CoA possibly derived from cellular protein valine. Concurrently a leucine related fatty acid increases significantly, potentially comprising the predominant pre-spore septum fatty acid component. These processes occur in conjunction with the onset of fatty acid β-oxidation and bulk protein turnover.

## Introduction

While the two fundamental biological processes of vegetative growth and cellular differentiation are separable only in principle, the sporulation process in *Bacillus subtilis* has long served as a model of the latter. The physiological, genetic, and regulatory aspects of this process have been the subject of intense research and are well documented in almost 8,000 literature reports. Changes in intermediary metabolism that are known or suspected to occur as growth slows and ceases are not as clearly understood, having only seldom been studied, either directly by means of quantitative metabolomics (Chubukov and Sauer, 2014), or indirectly using quantitative proteomics (Maaβ et al., 2014) or changes in gene expression (Koburger et al., 2005). The degradation of cellular fatty acids which occurs when CcpA-mediated carbon catabolite repression eases upon glucose depletion is probably the best understood aspect of early stationary-phase metabolism (Fujita et al., 2007; Matsuoka et al., 2007; Tojo et al., 2011). The other prominent metabolic process during sporulation, both selective and more generalized protein degradation, has also been intensively studied (Gerth et al., 2008; Kock et al., 2004; Schultz et al., 2017).

An intriguing feature of early stationary-phase metabolism that has received less research is its dependence on *de novo* biosynthesis of fatty acids, first reported by Schujman et al. (1998). These researchers found that such synthesis was required for activation of the mother cell-specific transcription factor σ^E^, suggesting that localization of this process might depend on asymmetric fatty acid composition of the pre-spore septal membrane. Martinez et al. (2010) showed that cells depleted in acyl carrier protein (ACP), essential for fatty acid synthesis, produced normal amounts of pro-σ^E^ but were unable to process it, confirming that *de novo* fatty acid synthesis was necessary for mother cell gene expression. Pedrido et al. (2013) found that *de novo* fatty acid synthesis was reactivated by transcription factor Spo0A only in the mother cell compartment, with maximal lipid synthesis occurring 2 to 4 hours after the onset of stationary phase, concurrent with asymmetric division of the cells. They also observed that *B. subtilis* required *de novo* fatty acid synthesis for biofilm formation, another Spo0A-dependent differentiation process (Lopez et al., 2009). Most recently, Haggett et al. (2018) detailed how Spo0A directly regulates expression of acetyl-CoA carboxylase, which catalyzes formation of malonyl-CoA in the first committed step of fatty acid biosynthesis.

Another intriguing feature of *Bacillus* metabolism is the centrality of the branched-chain amino acids (BCAAs), with the predominance of branched-chain fatty acids in the membranes of bacilli being the best-known feature of this relationship (Kaneda, 1977). Perhaps the most enigmatic property of the branched-chain amino acids is their control, along with GTP, of the activity of global regulatory protein CodY, which in turn participates in modulating their biosynthesis (Brinsmade et al., 2010). Indeed, “By mediating the creation of different steady-state levels of BCAA biosynthesis, under different conditions of nutrient availability, CcpA and CodY determine the extent to which CodY is active, thereby determining the impact of CodY on the many genes that it controls” (Sonenshein, 2007). Given that each operon or gene controlled by CodY is differentially regulated depending on the latter’s activity (Brinsmade et al., 2014), the degree of expression of these genes is apparently a function of the momentary concentration of isoleucine, valine and GTP.

Branched-chain amino acid metabolism also appears to have some direct yet undefined role in cellular differentiation of Firmicutes such as *Bacillus* and *Clostridium*, since the gene encoding the transcriptional activator of the branched-chain dehydrogenase enzymes, *bkdR*, is the most conserved gene amongst endospore-forming bacteria (Traag et al., 2013). Of particular interest in the present context are long neglected reports that *B. subtilis* metabolized radiolabeled valine into not only the expected “iso-even” branched-chain fatty acids but also into straight-chain (normal) and other types, incorporating label into the straight-chain segment of branched-chain fatty acids (Kaneda, 1963a,b). These results imply that either valine is metabolized to cellular intermediates like acetyl-CoA, or that a more direct pathway exists from valine into the backbone of fatty acids. While leucine has been shown to be capable of catabolism to acetyl-CoA by enzymes of the *yng* operon under σ^E^ control (Hsiao et al., 2010), it is unknown whether valine or isoleucine can be metabolized beyond their acyl-CoA esters which prime fatty acid synthesis.

In the following we report that portions of the carbon skeletons of valine and methionine were incorporated into the straight-chain segment of specific cellular fatty acids during late vegetative growth or “transition” phase in a complex sporulation medium. We also found a consistent change in cellular fatty acid composition during early stationary phase, congruent with the hypothesis of Schujman et al. (1998) that asymmetric fatty acid composition of the pre-spore septum might provide a mechanism for targeting σ^E^ activation to the mother cell compartment.

## Results

Unless stated otherwise, all results are for *Bacillus subtilis ΔtrpC2*, commonly known as the “168” strain (Zeigler et al., 2008), grown on complex sporulation medium. For those unfamiliar with the biochemistry of branched-chain amino and fatty acids, the biosynthetic pathway of the latter from the former is shown schematically in Figure 1.

**FIGURE 1.**
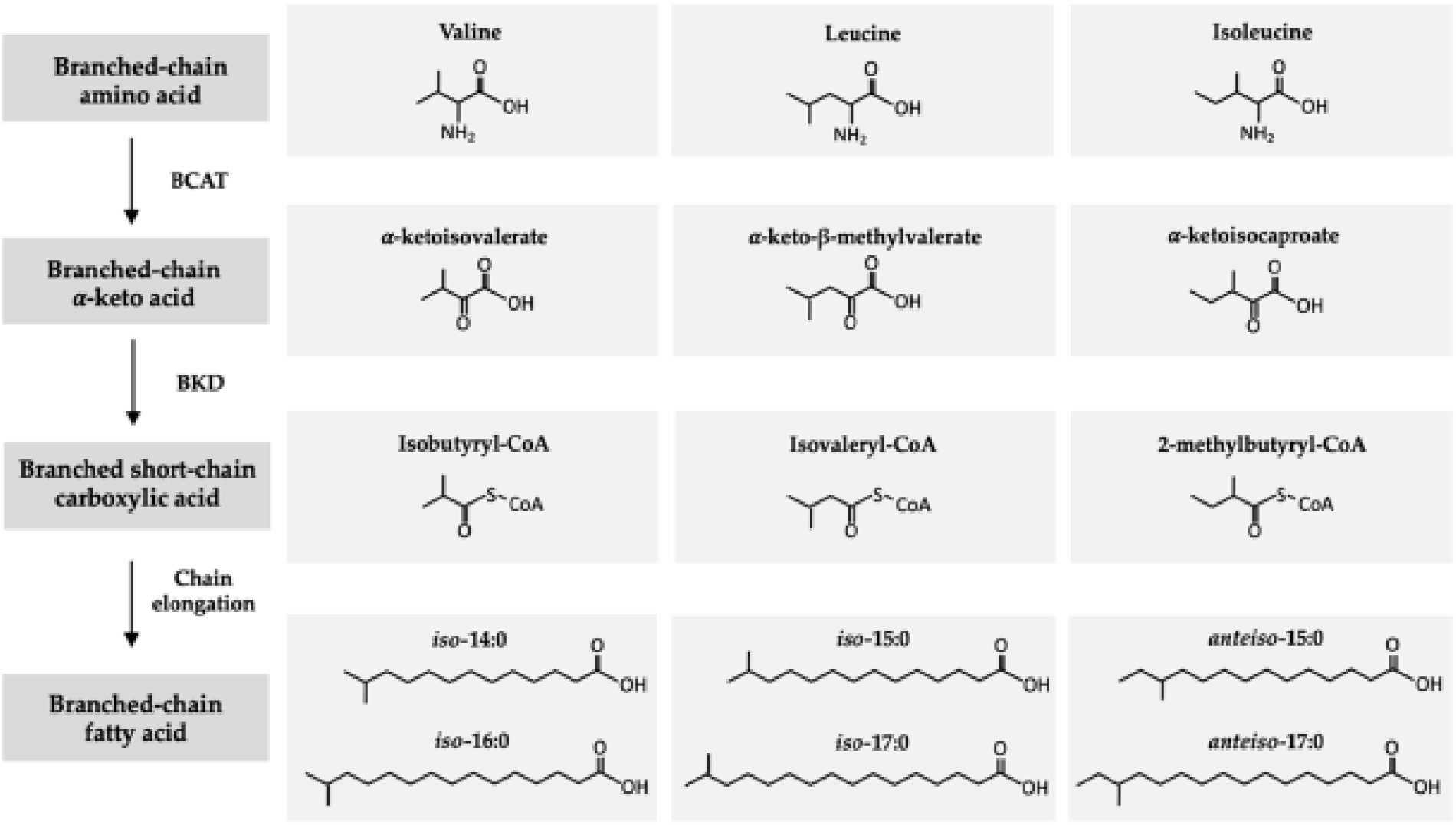
Biosynthetic pathway from branched-chain amino acids to branched-chain fatty acids. Abbreviations: BCAT: branched-chain amino acid transferase; BKD: branched-chain α-ketoacid dehydrogenase. Reproduced from Taormina et al. (2020) under the terms and conditions of the Creative Commons Attribution (CC BY) license (http://creativecommons.org/licenses/by/4.0/).

Since we will refer repeatedly to specific types of fatty acid molecule, the abbreviations that will be used for them are shown in Table 1, ordered according to the chromatographic retention time of their methyl esters. The abbreviation “i” is short for terminally iso-branched (*iso*-14:0 to *iso*-17:0 in Figure 1), “a” for terminally anteiso-branched (*anteiso*-15:0 and-17:0 in Figure 1), and “n” for normal (unbranched or straight-chain) fatty acids.

**TABLE 1.**
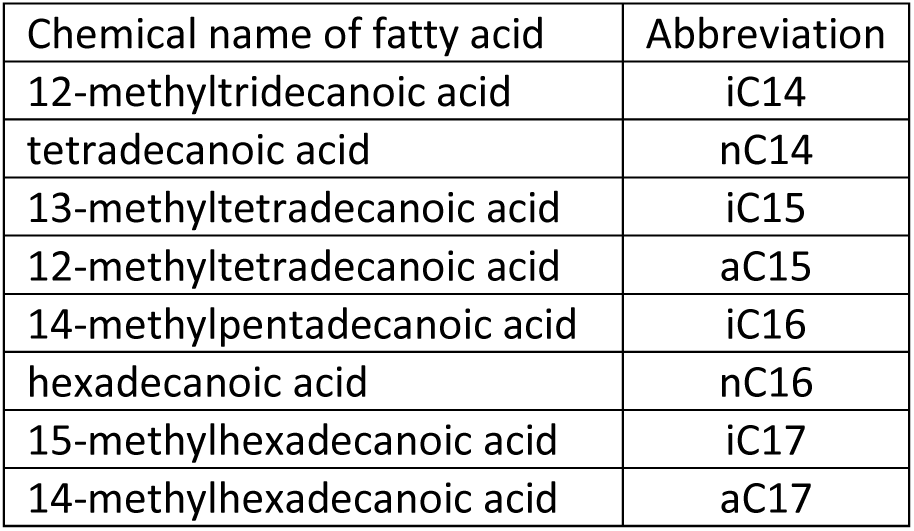
Summary of abbreviations used for fatty acids.

| Chemical name of fatty acid | Abbreviation |
| --- | --- |
| 12-methyltridecanoic acid | iC14 |
| tetradecanoic acid | nC14 |
| 13-methyltetradecanoic acid | iC15 |
| 12-methyltetradecanoic acid | aC15 |
| 14-methylpentadecanoic acid | iC16 |
| hexadecanoic acid | nC16 |
| 15-methylhexadecanoic acid | iC17 |
| 14-methylhexadecanoic acid | aC17 |

### Unexpected heavy analogs of iso-even fatty acids arise from deuterated valine

As fully described in Experimental Procedures, we used capillary gas chromatography coupled to mass spectrometry to analyze *B. subtilis* cellular fatty acids in the hours following vegetative growth on sporulation medium supplemented with deuterium labeled amino acids. More specifically, at hourly intervals between 3 and 6 hours of culture, we scanned the mass-to-charge (*m/z*) range of 60 to 360 for the eluting methyl ester peaks of all expected fatty acids (Kaneda, 1977). This allowed us to identify most peaks by comparison of their mass spectra to standard databases and their relative retention times, as well as by injection of *n*-tridecanoic acid, nC14 and iC16 standards. Two apparently anomalous peaks were present, however, eluting just prior to iC14 and iC16 in cultures supplemented with *L*-valine-d_8_ but not in those supplemented with unlabeled *L*-valine, as shown in Figure A1(a) to A1(d). Since the “iso-even” fatty acids iC14 and iC16 are synthesized using as primer isobutyryl-CoA (Figure 1), and since much of the pool of this primer consisted of isobutyryl-d_7_-CoA when *L*-valine-d_8_ was added to the medium (see below), we suspected that the unknown peaks were primed with the deuterated ester. This suspicion was supported by the mass spectra of the various peaks which implied that the “mother” ion (M) of iC14 at approximately *m/z* 242 (Figure A1(c)) had shifted to approximately *m/z* 249 in the earlier eluting peak (Figure A1(a)), and that of iC16 had shifted from approximately *m/z* 270 (Figure A1(d)) to approximately *m/z* 277 in the earlier eluting peak (Figure A1(b)). Similar 7 Da increases in mass were also evident in several fragments arising from the early eluting peaks. Based on this evidence we concluded that these “anomalous” peaks represented fatty acids primed with isobutyryl-d_7_-CoA and hence were perdeuterated in the terminal three carbon atoms; herein they are designated “iC14-d7” and “iC16-d7” to distinguish them from iC14 and iC16 formed with unlabeled isobutyryl-CoA as primer.

Using an *m/z* 60-360 scan we initially measured the intensity of the singly charged (M-1) to (M+9) ions to detect any heavy analogs (mass isotopomers) of each fatty acid present. To avoid confusion, for the fatty acids designated iC14-d7 and iC16-d7 we will designate in the text the “mother” ions with *m/*z values of approximately 249 and 277, respectively, with the notation “M_d7_”; so, for these we recorded the intensity of the (M_d7_-1) to (M_d7_+9) ions. We corrected each experimental isotopomer distribution for loss of hydrogen from the mother ion and/or tailing of M into the (M-1) peak (González-Antuña et al., 2014; ditto for M_d7_), and finally calculated the enrichment, if any, of each analog relative to natural abundance via the commonly used correction matrix method (Midani et al., 2017; Nanchen et al., 2007). These results for cultures of *B. subtilis ΔtrpC2* (168) and *Δspo0A::ermtrpC2* strains supplemented with *L*-valine-d_8_ gave apparent isotopic enrichment of a number of heavy analogs in several fatty acids. However, since the isotopomer distributions after correction for natural abundance gave one or more negative values for most fatty acids, indicative of inaccuracy in the data (Midani et al., 2017), we decided to measure the intensity of only the (M-1) to (M+5) ions [or (M_d7_-1) to (M_d7_+5) ions] for each fatty acid, and to do so during a period encompassing its chromatographic retention time, as described in detail in Experimental Procedures. The primary mass spectrometric data files for triplicate cultures supplemented with *L*-valine-d_8_ (RC129, RC130 and RC131), and triplicate cultures supplemented with both *L*-valine-d_8_ and *D*,*L*-methionine-d_4_ (RC128, RC132 and RC133), are appended as “Rowe & Perreault spreadsheets”.

This methodology, when applied to cultures of the *ΔtrpC2* strain supplemented with *L*-valine-d_8_, produced corrected distributions with very few negative values due to missing or low intensity peaks (“Val-d8 enrichment calculations.xlsx”). These results showed by far the strongest enrichment of an (M_d7_+4) analog (or analogs), present only in iC14-d7 and iC16-d7 fatty acids potentially derived from protein breakdown. The composite chromatographic peak for these fatty acids, an example of which is shown in Figure A2, was partially resolved into the expected iC16-d7 species with an *m/z* value for the mother ion (M_d7_) of approximately 277, and a trailing peak with a corresponding *m/z* value of approximately 281, evidently containing an additional four deuterium atoms. Given that the branched portion of the mother ion is completely deuterated, the additional deuterium could only be appended to carbon atoms in the straight-chain fatty acid “backbone”. The signal-to-noise ratio of these two peaks ranged from 8 to 9, giving an estimated coefficient of variation (CV) for the signal of approximately 6 % (Dolan, 2006). A similar enrichment of the (M_d7_+4) analog of iC14-d7 was also observed in triplicate *B. subtilis ΔtrpC2* cultures supplemented with *L*-valine-d_8_, although the CV of the 3-hour enrichment of this analog in the iC14-d7 pool was about 11 %, while the CV of the (M_d7_+4) analog in the iC16-d7 pool was about 1.5 %. (“Val-d8 isotopomer enrichment calculations.xlsx”).

Figure 2 shows that both iC14-d7 and iC16-d7 were significantly (*P* < 0.01) enriched in an (M_d7_+4) analog throughout the 3-to-6-hour period studied, with the highest enrichment appearing to occur from 3-4 hours. To simplify the notation, the (M_d7_+4) mass isotopomer is alternatively designated as “iC14-d11” or “iC16-d11”, and the (M_d7_+5) analog designated as “iC14-d12” or “iC16-d12”.

**FIGURE 2.**
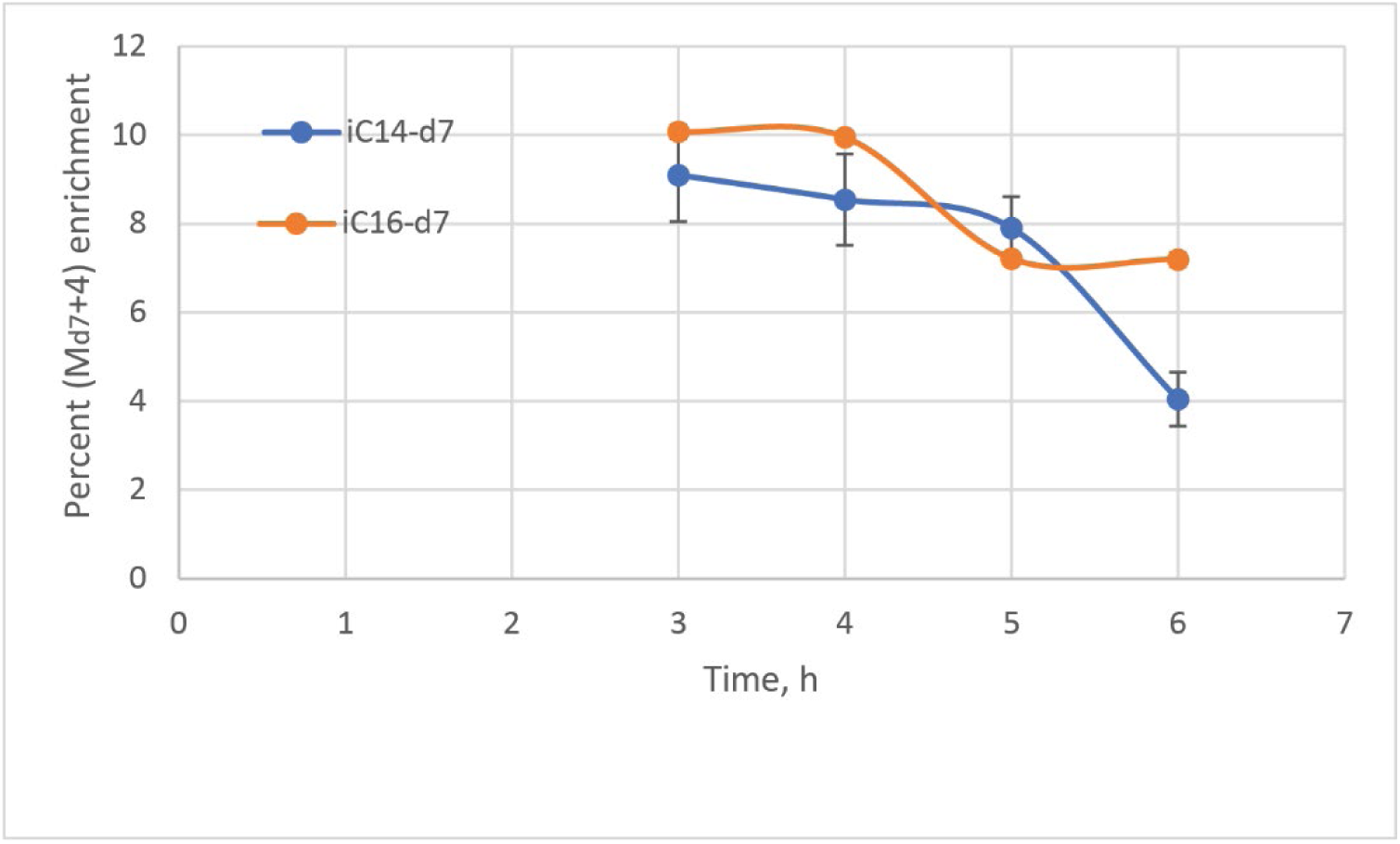
Temporal profile of percent (M_d7_+4) enrichment of iC14-d7 and iC16-d7 fatty acids in triplicate *B. subtilis ΔtrpC2* sporulation medium cultures supplemented with *L*-valine-d_8_. Data points are the percentage of iC14-d11 in the iC14-d7 pool, and iC16-d11 in the iC16-d7 pool. Error bars represent one standard error of the mean (SEM) of replicate cultures (n=3); the SEM values for iC16-d7 of ≤ 0.16 % are too small to be visible.

That this (M_d7_+4) analog resulted from *B. subtilis* metabolism rather than from an analytical artifact was verified by its absence from the strong iC14-d7 and iC16-d7 peaks of a *Xanthomonas campestris* culture supplemented with 2-methylpropionic-d_7_ acid, a direct precursor of isobutyryl-d_7_-CoA. As Figure A3(b) shows, the *m/z* 253 peak characteristic of the (M_d7_+4) isotopomer of iC14-d7 was virtually absent; similarly for the *m/z* 281 peak of the (M_d7_+4) analog of iC16-d7 in Figure A3(d). The apparent enrichment of these analogs was no more than 0.3 ± 0.1 % (“X_campestris enrichment calculations.xlsx”), thus showing that the 8-10 % enrichment observed in *B. subtilis* (Figure 2) was the result of metabolism of *L*-valine-d_8_.

### Fatty acid beta-oxidation produces lower molecular weight analogs

In *B. subtilis ΔtrpC2* sporulation-medium cultures supplemented with *L*-valine-d_8_, enrichment of (M+1) and (M+2) analogs of most fatty acids, significant at the *P* < 0.01 level, was observed throughout the period studied (“Valine-d8 enrichment calculations.xlsx”). For the most prominent fatty acid until at least 4 hours (aC15), (M+1) and (M+2) enrichment averaged 2.68 (SEM = 0.09) and 0.91 per cent (SEM = 0.07), respectively (both n = 12). In all cases where there was significant enrichment of both (M+1) and (M+2) analogs, the amount of the latter was only a fraction of that of the former, implying that both were likely formed by incorporation of units of acetyl-d-CoA into the fatty acid via malonyl-d-CoA/-ACP. This labeling scenario is to be expected from the sequence of β-oxidation of a fatty acid having a deuterated backbone followed by fatty acid biosynthesis using the acetyl-d-CoA so derived (Fujita et al., 2007).

Given that increases in water deuteration are a standard method for measuring the fatty acid oxidation rate (Ecker & Liebisch, 2014; Law et al., 2007), we used isotope ratio mass spectrometry to study culture broth water composition versus time. As shown in Figure A4, substantial deuteration of broth water occurred in cultures of the *B. subtilis ΔtrpC2* strain supplemented with *L*-valine-d_8_. Based on these results we estimate that 7.1 % of the deuterium in the valine added was oxidized between 3 and 6 hours, consistent with induction of the enzymes responsible for β-oxidation upon easing of carbon catabolite repression (Fujita et al., 2007). Analysis of single *B. subtilis* 168 trp+ cells has shown that upon glucose starvation transcription of two key β-oxidation genes, *fadE* and *fadN*, is up-regulated 92-and 74-fold, respectively (de Jong et al., 2012, their Table A4).

### Unexpected heavy analogs of iso-even fatty acids also arise from valine-^13^C_5_

In early work using the *m/*z 60-360 scan method, we studied fatty acid analogs in triplicate cultures of *B. subtilis ΔtrpC2* grown on sporulation medium supplemented with *L*-valine-^13^C_5_,^15^N, the results of which are shown in the appended file “Val-13C5 isotopomer data.xlsx”. Not unexpectedly, enrichment of 8 to 9 per cent was found in the (M+4) isotopomer of iC14 and iC16 fatty acids, evidently primed with isobutyryl-^13^C_4_-CoA (“Val-13C5 isotopomer enrichment calculations. xlsx”). What was not expected, and whose significance was recognized only later, was the generally significant (*P* < 0.01) presence of (M+6) analogs in these fatty acids. As discussed below, the fact that no (M_d7_+6) isotopomers were detectable in similar cultures supplemented with *L*-valine-d_8_ (“Val-d8 isotopomer enrichment calculations.xlsx”) implied that either two carbon-13 atoms had been incorporated into the straight-chain segment of the iso-even fatty acids primed with isobutyryl-^13^C_4_-CoA, or six carbon-13 atoms had been incorporated into the straight-chain segment of the iso-even fatty acids primed with unlabeled isobutyryl-CoA.

### Acyl-CoA concentrations change during early sporulation phase

Because of the prominent role of acyl-CoA esters in branched-chain amino and fatty acid metabolism (Figure 1), we studied the intracellular concentration of the relevant branched-chain CoA esters of the *ΔtrpC2* strain under sporulation and non-sporulation conditions. Given the distinct possibility that isobutyryl-CoA might be further metabolized to propionyl-CoA, research on which is summarized in Appendix 2, analysis of this ester was also performed. As shown in Table 2 for cultures supplemented with *L*-valine-d_8_, in the absence of early sporulation-phase metabolism propionyl-CoA accumulated markedly as an apparently unmetabolizable product derived from isobutyryl-CoA, whereas the former ester was almost undetectable a couple of hours after the transition to sporulation metabolism began.

**TABLE 2.** Intracellular acyl-CoA ester concentration (μM) at 6-7 hours for triplicate B. subtilis ΔtrpC2 cultures on non-sporulation and sporulation medium supplemented with L-valine-d_8_.

| Acyl-CoA | Non-sporulation |  | Sporulation |  |
| --- | --- | --- | --- | --- |
|  | Mean | SEM<sup>a</sup> | Mean | SEM<sup>a</sup> |
| isobutyryl- | 2.8 | 0.5 | 14.3 | 3.2 |
| isobutyryl-$d_7$- | 5.7 | 0.8 | 24.6 | 6.9 |
| propionyl- | 52.0 | 24.8 | 0.2 | 0.2 |
| propionyl-$d_5$- | 18.5 | 7.6 | 0.0 | 0.0 |

<sup>a</sup> SEM of replicate cultures, n=3

Figure 3 shows the temporal evolution of deuterated and non-deuterated isobutyryl-and propionyl-CoA in sporulation medium cultures supplemented with *L*-valine-d_8_. Although with substantial variability from culture to culture, the concentration of both labeled and unlabeled isobutyryl-CoA appeared to increase during early sporulation, while the little propionyl-CoA initially present vanished by 4 hours.

**FIGURE 3.**
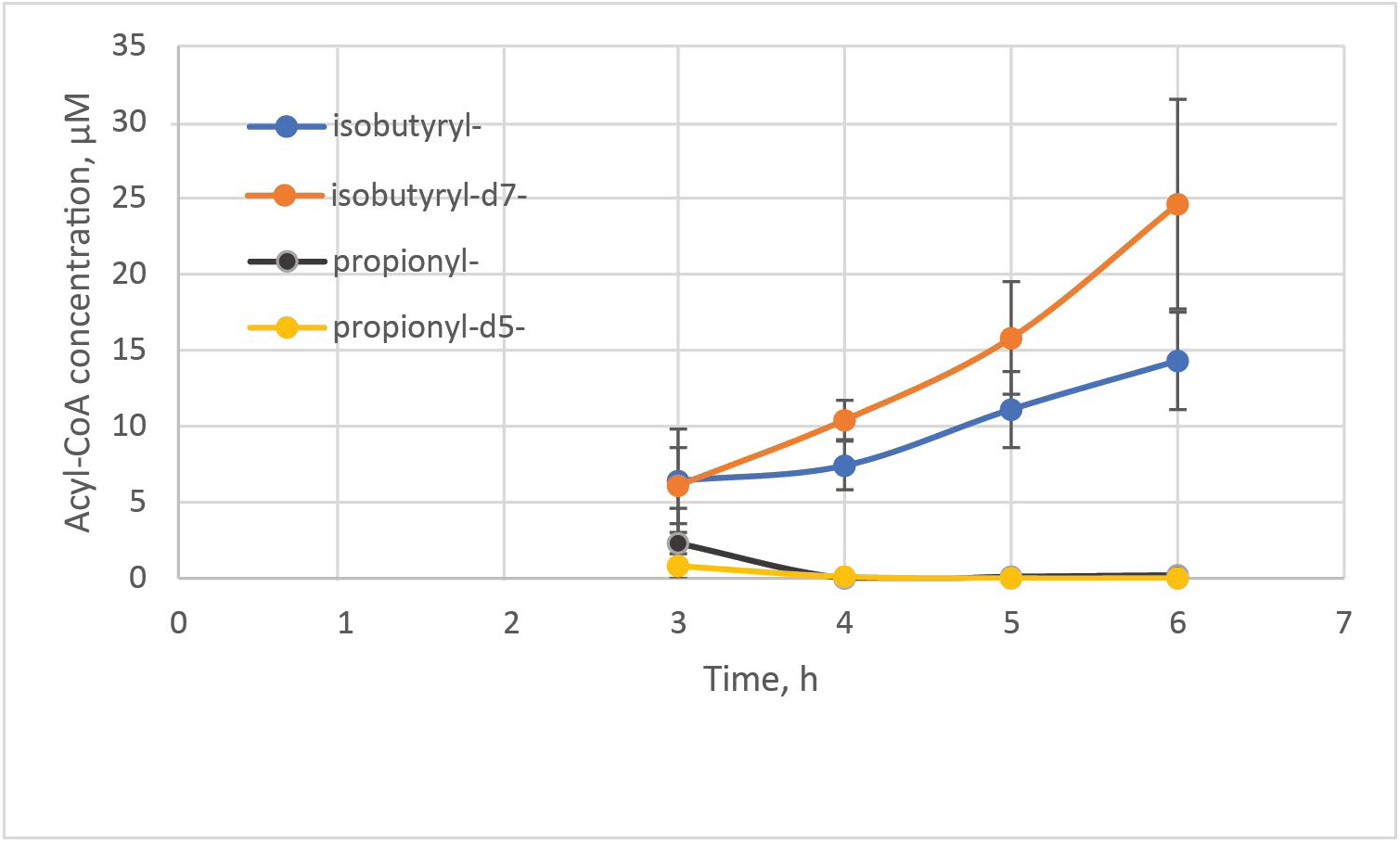
Temporal profile of isobutyryl-CoA and propionyl-CoA concentrations in triplicate *B. subtilis ΔtrpC2* sporulation medium cultures supplemented with *L*-valine-d_8_. Error bars represent one SEM of replicate cultures (n=3).

Having established that *B. subtilis* metabolizes valine via isobutyryl-CoA to propionyl-CoA, it was of interest to know if the latter ester could also be produced from isoleucine (Massey et al., 1976). However, *B. subtilis ΔtrpC2* grown on three different media supplemented with *L*-isoleucine-d_10_ was found to produce abundant 2-methylbutanoyl-d_9_-CoA but did not process it further to the expected metabolite, propionyl-d_5_-CoA. Most tellingly, in cells grown in non-sporulation medium, which gave substantial propionyl-d_5_-CoA from *L*-valine-d_8_ (Table 2), no labeling of this ester was detectable from *L*-isoleucine-d_10_.

### Iso-even fatty acid analogs are also formed from deuterated methionine

Since the acyl-CoA results led us to suspect propionyl-CoA as an intermediate in the evident pathway from valine to the straight-chain segment of iso-even fatty acids, we studied cultures supplemented with another amino acid capable of being metabolized to propionyl-CoA, namely *D*,*L*-methionine. Conversion of *L*-methionine to cysteine forms 2-oxobutanoate as a by-product (Hullo et al., 2007), which both the branched-chain 2-oxoacid dehydrogenase complex (BkdAA/AB/B) and pyruvate dehydrogenase complex (PDH) can convert to propionyl-CoA (Oku & Kaneda, 1988). *B. subtilis* has also been shown to use *D*-methionine for growth (Hullo et al., 2004), apparently by converting it to the *L*-isomer.

As expected, cultures of *B. subtilis ΔtrpC2* on sporulation medium supplemented with a combination of *L*-valine-d_8_ and *D*,*L*-methionine-d_4_ produced iC14-d7 and iC16-d7 fatty acids significantly enriched in an (M_d7_+4) mass isotopomer (“Val-d8+Met-d4 isotopomer enrichment calculations.xlsx”). As Figure 4(a) shows, at 3 hours the amount of iC16-d11 plus iC16-d12 present was significantly higher (*P* < 0.01) when both *D*,*L*-methionine-d_4_ and *L*-valine-d_8_ were added compared to the case when the latter was added alone or in combination with unlabeled methionine. This result demonstrated that carbon from both methionine and valine was incorporated into iC16 fatty acid molecules primed with isobutyryl-d_7_-CoA originating from *L*-valine-d_8_. That iC16-d11 plus iC16-d12 was lower at 5 hours in the cultures supplemented with both *D*,*L*-methionine-d_4_ and *L*-valine-d_8_ is addressed in the Discussion.

**FIGURE 4.**
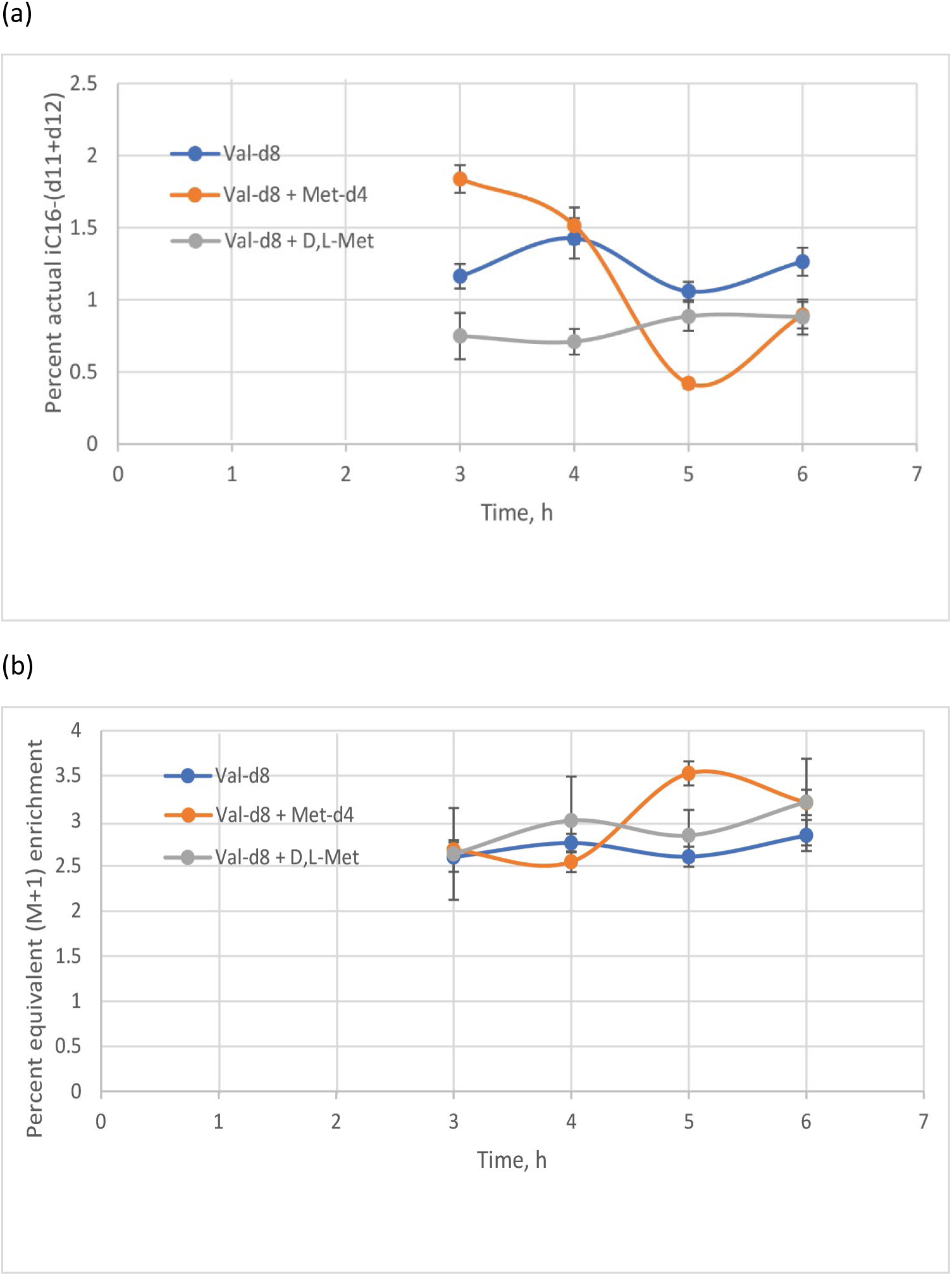
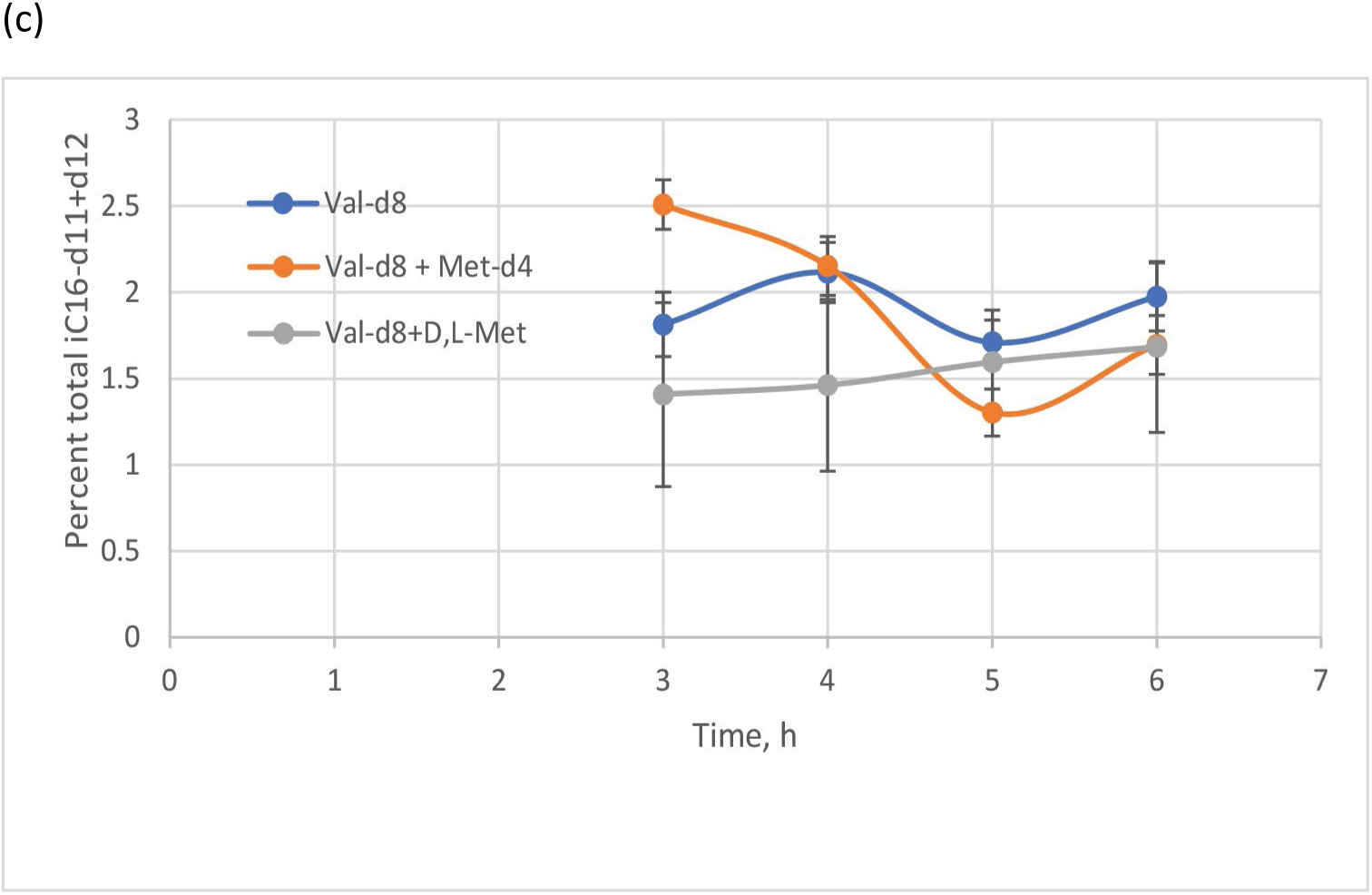
Temporal profile of (a) iC16-d11 plus iC16-d12 fatty acids; (b) the weighted equivalent (M+1) enrichment of all fatty acids; (c) iC16-d11 plus iC16-d12 fatty acids, including those lost to β-oxidation; all as a percentage of total cellular fatty acids, in triplicate *B. subtilis ΔtrpC2* sporulation medium cultures supplemented with *L*-valine-d_8_, *L*-valine-d_8_ and *D*,*L*-methionine-d_4_, or *L*-valine-d_8_ and *D*,*L*-methionine. Error bars represent one SEM of replicate cultures (n=3).

Figure 4(b) shows the extent of deuteration of cellular fatty acids versus time due to incorporation of acetyl-d-CoA (or ACP-d-CoA; Srinivas, 2018), more explicitly the sum of the fractional enrichment of (M+1) and (M_d7_+5) analogs, plus twice that of the (M+2) analogs (assumed to arise from incorporation of two acetyl-d-CoA molecules), weighted by the percentage of each fatty acid relative to total fatty acids. At 5 hours, for reasons that are not apparent, weighted equivalent (M+1) fatty acid enrichment was significantly higher and iC16-d11 plus iC16-d12 significantly lower (Figure 4(a); both *P* < 0.01) in the cultures supplemented with both *L*-valine-d_8_ and *D*,*L*-methionine-d_4_ compared to those supplemented only with *L*-valine-d_8_.

To estimate the activity of the pathway from valine and methionine into the observed (M_d7_+4) and (M_d7_+5) analogs we combined the data of Figures 4(a) and (b) to give the amount of iC16-(d11 + d12) that would be present in the absence of β-oxidation. These results, shown in Figure 4(c), again demonstrate that at 3 hours the amount of deuterated analogs present was significantly higher (*P* < 0.05) when the cultures were supplemented with both *D*,*L*-methionine-d_4_ and *L*-valine-d_8_ compared to the case when only the latter was present. The variability of results with the *L*-valine-d_8_ plus *D*,*L*-methionine cultures prevents any corresponding inference. Interestingly, there was no significant difference after 3 hours between the Val-d8 and Val-d8 plus Met-d4 cultures in the total amount of iC16-(d11 + d12) estimated to be present if β-oxidation were not occurring.

### Cellular fatty acid composition changes during B. subtilis differentiation

As shown in Figure 5 for the nine cultures reported above for which we obtained segmented-scan GC-MS data, overall mean values for anteiso-branched and normal species decreased as a percentage of total fatty acids, and their iso-branched counterparts increased essentially monotonically after 3 hours (cf. Table A1). A significant (*P* < 0.01) 3-6 hour increase in iC15 was especially evident. In fact, the increase in total iso-branched fatty acids was due entirely to iC15, since there was no significant change in iso-even types, iC17 fatty acids generally decreased over this period (Table A2), and *B. subtilis* does not produce iC13 or shorter fatty acids (Kaneda, 1977). We describe this phenomenon as characteristic of *B. subtilis* differentiation, rather than sporulation *per se*, because it also occurred in cultures of the *Δspo0A::ermtrpC2* strain grown under the same conditions (Figure A5, Table A1).

**FIGURE 5.**
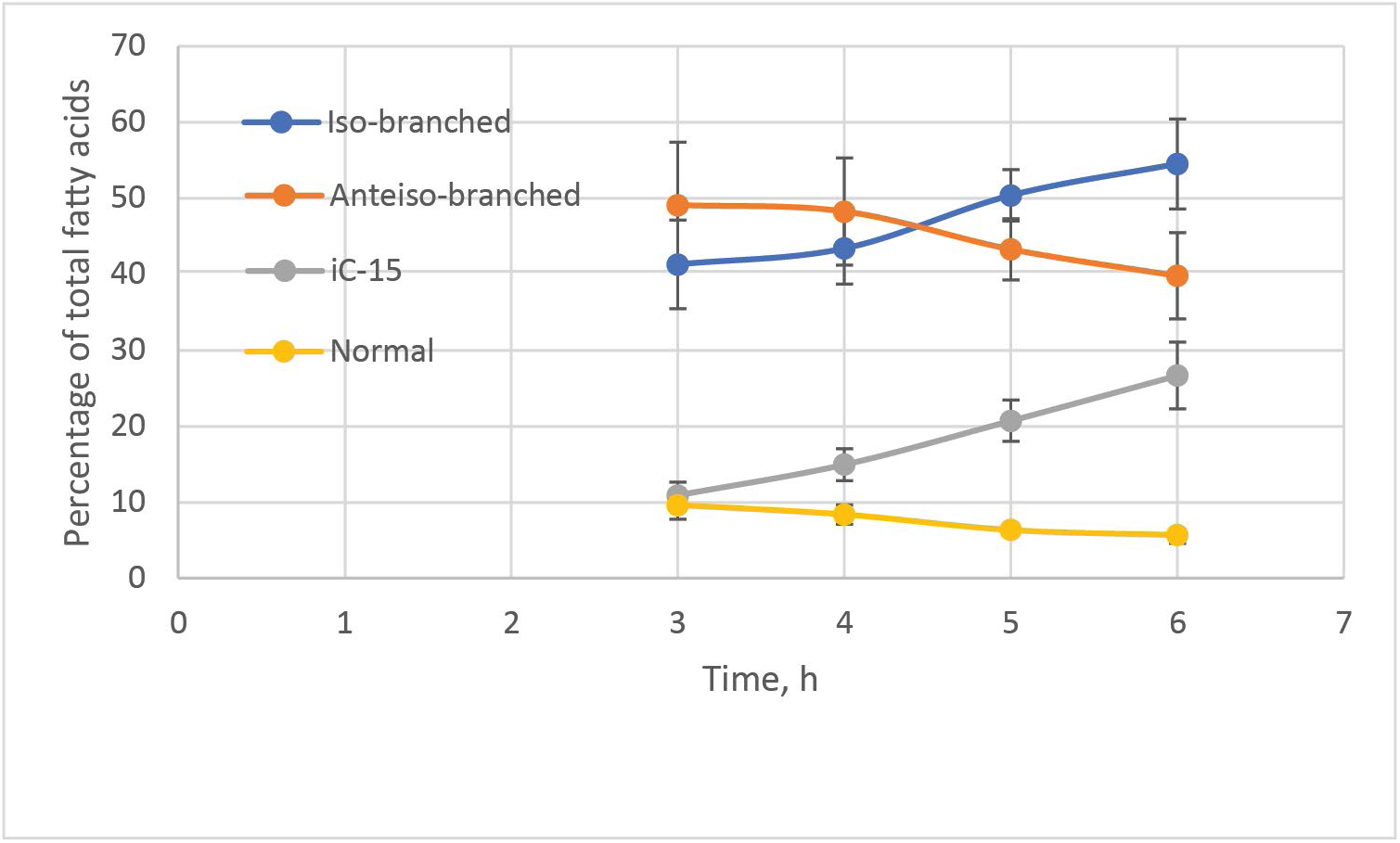
Temporal profile of *B. subtilis ΔtrpC2* iso-branched, anteiso-branched, iC15 and normal fatty acids in a total of nine sporulation medium cultures supplemented with *L*-valine-d_8_, *L*-valine-d_8_ and *D*,*L*-methionine-d_4_, or *L*-valine-d_8_ and *D*,*L*-methionine. Error bars represent one overall SEM for triplicate cultures and each of the three supplementation treatments (n=9).

## Discussion

To make the metabolic phenomena to be discussed easier to envisage, a schematic view of the apparent composite pathway derived from analysis of the results is shown in Figure 6. As indicated by the color coding in the Figure, it transforms part of the carbon skeleton of valine and methionine into a segment of the backbone of iso-even branched-chain fatty acids made exclusively using isobutyryl-CoA possibly formed from protein-derived valine as primer.

**FIGURE 6.**
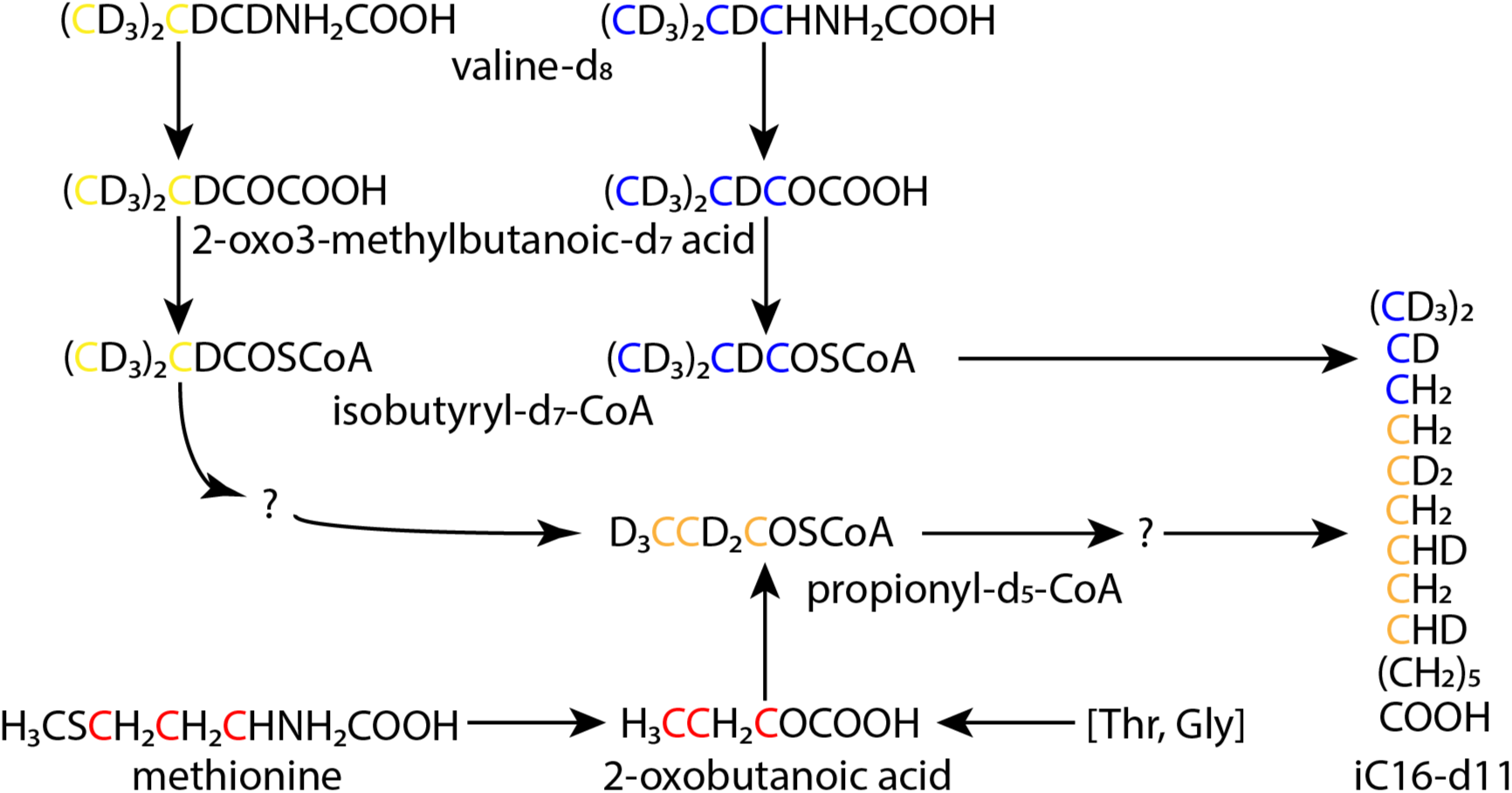
Schematic view of the apparent composite pathway from valine and methionine to iC16 fatty acid. Carbon atoms in orange originate from both valine (yellow) and methionine (red). For clarity, deuterium atoms originating specifically from methionine are not shown. Potential inputs are shown in square brackets. The location within the iC16 straight-chain segment of the carbon and deuterium atoms coming from propionyl-CoA is uncertain.

### Characteristics of the pathway from valine to fatty acids

Firstly, we note that the mass spectrum of the leading iC16-d7 fatty acid peak shown in Figure A2 (b) is proof of the presence of its (M_d7_+4) analog, i.e., an iC16-d11 species, since the (M_d7_+5)/(M_d7_+4) intensity ratio is essentially identical to that of (M_d7_+1)/M_d7_. This strongly implies that the analog has the same carbon skeleton as the mother ion, sharing identical amounts of natural-abundance carbon-13 and deuterium arising from acetyl-d-CoA. That the (M_d7_+4) analog is not an artifact of electron-impact ionization is proven by its essential absence from the iC14-d7 and iC16-d7 mass spectra of *X. campestris* (Figure A3). We also note that, although significant enrichment of (M_d7_+4) and (M_d7_+5) analogs was found only in fatty acids primed with isobutyryl-d_7_-CoA, it is likely that a small amount primed with unlabeled isobutyryl-CoA was present but undetected, since only 69.8 ± 0.2 % of total cellular valine was labeled in *B. subtilis ΔtrpC2* cells grown 9 hours on sporulation medium supplemented with *L*-valine-^13^C_5_.

Our results with the *B. subtilis ΔtrpC2* strain in sporulation medium (Figure 2) show that the valine to fatty acid pathway is induced by 3 hours, i.e., at least as early as the slowing or cessation of vegetative growth. That exponential growth had ended by 3 hours is shown by the essentially linear rate of biomass increase for at least two hours thereafter (Figure A6). Although we cannot rule out the possibility that the pathway also operates during growth phase, we hypothesize that it is more likely associated with cellular differentiation since it appears to be operative substantially contemporaneously with cellular protein turnover and fatty acid β-oxidation. Whether the valine and methionine that enter this pathway are necessarily derived from cellular protein breakdown will be the subject of future investigation.

We found evidence that this pathway is associated with cellular differentiation rather than sporulation *per se* since the *B. subtilis Δspo0A::ermtrpC2* strain grown under the same conditions gave 3-hour (M_d7_+4) enrichment of iC14-d7 even greater than that of the *ΔtrpC2* strain (*P* < 0.01; Table A3). Analysis of intracellular acyl-coenzyme A esters showed not only that *B. subtilis* metabolizes valine via isobutyryl-CoA to propionyl-CoA, but that the relative concentration of these metabolites differs markedly between sporulating and non-sporulating cells (Table 2). Accumulation of propionyl-CoA in the latter, and its virtual disappearance in the former (Figure 3), implies that it is further metabolized during the transition from growth to sporulation.

The identity of the enzymes that convert isobutyryl-CoA to propionyl-CoA has not been definitively established, and a summary of work on this question and remaining uncertainties is given in Appendix 2. As noted there, experiments with *B. subtilis* strains lacking specific enzymes indicate some level of redundancy in the pathway from valine to propionyl-CoA.

The results of cultures supplemented with both *L*-valine-d_8_ and *D*,*L*-methionine-d_4_ (Figure 4) proved that the pathway into the straight-chain segment of iso-even fatty acids passes through propionyl-CoA, as shown in Figure 6. In addition to the strong (M_d7_+4) labeling observed with these deuterated amino acids (Figure 2), the much weaker but apparently significant presence of (M+6) analogs was observed in cultures supplemented with *L*-valine-^13^C_5_,^15^N (“Val-13C5 isotopomer enrichment calculations. xlsx”). To substantiate that this latter observation is real it is necessary to rationalize why the calculated carbon-13 enrichment of iC16 might be at least an order of magnitude lower than that with deuterium labeling.

The first factor to consider is that capillary GC separated fatty acids primed with isobutyryl-d_7_-CoA from those primed with unlabeled isobutyryl-CoA, whereas no such separation occurred when isobutyryl-^13^C_4_-CoA was the primer. Thus, assuming that the cells do not metabolically differentiate between *L*-valine-d_8_ and *L*-valine-^13^C_5_,^15^N (i.e., absent any kinetic isotope effect), in the latter case the iC16 fatty acids were primed with either isobutyryl-^12^C_4_-CoA or isobutyryl-^13^C_4_-CoA, and the precursor for incorporation into the fatty acid backbone was either propionyl-^12^C_3_-CoA or propionyl-^13^C_3_-CoA; in brief, four different sets of precursors can be identified in principle. Assuming no isotope effect on any relevant enzyme, to a first approximation the fraction of iC16 derived from *L*-valine-^13^C_5_,^15^N (let’s designate it “F”) can be calculated as the amount of iC16-d7 divided by the total of iC16-d7 plus the iC16 primed with isobutyryl-CoA. The second factor to consider is that in sporulation medium supplemented with *L*-valine-^13^C_5_,^15^N only approximately 70 % of total cellular valine was labeled; thus, in general presumably about 30 % was unlabeled by either carbon-13 or deuterium. Therefore, a more accurate estimate of the fraction of iC16 derived from *L*-valine-^13^C_5_,^15^N is given by (100/70) x F. Since iC16-d7 ranged from about 12-22 % of the total (iC16-d7 + iC16) pool (Table A2), the percentage of the corresponding pool originating from *L*-valine-^13^C_5_,^15^N was approximately 17-31 %.

Finally, there are two possible routes to the (M+6) analog detected in the iC16 fatty acids formed from *L*-valine-^13^C_5_,^15^N, namely: they were primed with isobutyryl-^13^C_4_-CoA and two carbon-13 atoms were subsequently inserted into the backbone; or they were primed with isobutyryl-^12^C_4_-CoA and six carbon-13 atoms were subsequently inserted into the backbone. That approximately 70 % of total cellular valine was labeled allows us to calculate the probability of occurrence for each route. For the former, labeled primer, route it is (0.70 x 0.70) ≈ 0.5; for the latter it is (0.30 x 0.70) ≈ 0.2. Thus, if an (M+2) analog were formed with isobtyryl-^13^C_4_ as primer the expected enrichment of an (M+6) analog would be 0.5 x 17-31 %, i.e., in the range of 9-16 % relative to that observed for deuterium labeling. Conversely if an (M+6) analog were formed with unlabeled isobutyryl-CoA as primer the expected enrichment would be 0.2 x 17-31 %, or in the range of 3-6 % relative to that observed for deuterium labeling. Since the observed enrichment of the (M_d7_+4) analog of iC16-d7 was on the order of 10 % (Figure 2), the two possible carbon-13 scenarios of (M+2) or (M+6) analog formation would give expected enrichments in the range of 0.9-1.6 % and 0.3-0.6 %, respectively. Hence, since the observed significant (P < 0.01) iC16 carbon-13 enrichments were in the range of 0.4-0.7 % (“Val-13C5 enrichment calculations.xlsx”), we are led to the tentative conclusion that an (M+6) isotopomer(s) was somehow formed from propionyl-^13^C_3_-CoA, as shown schematically in Figure 6.

In summary, our analysis of the results of labeling experiments leads to the conclusion that *B. subtilis* incorporated four deuterium atoms arising from propionyl-d_5_-CoA or six carbon-13 atoms arising from propionyl-^13^C_3_-CoA into the straight-chain backbone of iso-even fatty acids. Even if the carbon-13 data is for some reason erroneous, and propionyl-CoA could somehow be converted into malonyl-CoA or succinyl-CoA, and one or the other of these incorporated into a fatty acid chain, it could add only two deuterium atoms or would add three carbon atoms, respectively, both of which would have to occur exactly twice to produce the required labeled 14-or 16-carbon molecule. Another possibility might be condensation of two molecules of propionyl-CoA to 2-methyl-2-pentenoyl-CoA, as has been reported to occur in several types of mammalian cells (Doan et al., 2019; Snyder et al., 2015). This would require the activity of a mutase enzyme which has not been identified in *B. subtilis*, however, and we have been unable to construct a satisfactory pathway via 2-methyl-2-pentenoyl-CoA. Although completely speculative, we have synthesized a possible pathway from propionyl-CoA to fatty acids which could explain both the deuterium and carbon-13 labeling results, as described in detail in Appendix 3. It may not be coincidental that the hypothetical pathway includes many enzymes constitutive for fatty acid biosynthesis and β-oxidation, and three others that are strongly induced at t_2_ of sporulation.

### Significance of the pathway from methionine to fatty acids

Our 3-hour results with cultures of *B. subtilis ΔtrpC2* supplemented with *D*,*L*-methionine-d_4_ in addition to *L*-valine-d_8_ (Figure 4(a)) demonstrate with a high degree of confidence (*P* < 0.01) that carbon atoms of methionine are incorporated into the backbone of iso-even fatty acids. That there was no significant difference after 3 hours between these treatments in the amount of iC16-(d11 + d12) that would be present in the absence of β-oxidation (Figure 4(c)) is intriguing, and mighty imply that methionine catabolism to propionyl-CoA occurs only early in the differentiation process.

Although methionine carbon is known to pass into propionyl-CoA via incorporation into 2-oxobutanoate (Hullo et al., 2007) followed by the action of pyruvate dehydrogenase and/or branched-chain 2-oxoacid dehydrogenase (Oku & Kaneda, 1988), there is a prevalent misconception that methionine can serve as a carbon source for *B. subtilis*. This would presumably occur by conversion of propionyl-CoA to succinyl-CoA and entry of the latter into the TCA cycle, however this pathway appears not to be functional in this organism due to the absence of methylmalonyl-CoA mutase activity (Birch et al., 1993).

That 2-oxobutanoate (α-ketobutyrate) is an intermediate in the pathway from methionine (and potentially threonine and glycine) into fatty acids is intriguing since it is the essential precursor for isoleucine biosynthesis (Mӓder et al., 2004). Isoleucine is the most potent ligand affecting the activity of global regulatory protein CodY, especially during the transition from vegetative growth to stationary phase (Shivers & Sonenshein, 2004), hence its concentration indirectly regulates the expression of hundreds of stationary phase genes (Sonenshein, 2005). It would be tempting to hypothesize that induction of the pathway from valine and methionine to iso-even fatty acids depletes intracellular 2-oxobutanoate and reduces isoleucine concentration by preventing its biosynthesis, thereby inactivating CodY-mediated gene repression.

### Changes in cellular fatty acid content and the establishment of cell fate

Our results show that cellular fatty acid composition changed essentially monotonically from 3-6 h, with anteiso-branched and normal species decreasing, and a corresponding increase in iso-branched species driven entirely by increased iC15 (Figure 5, Table A2). Except for supplementation with 0.5 mg ml^-1^ of valine and methionine, the culture conditions that we employed were exactly those of Leighton and Doi (1971), who observed an exponential increase in alkaline phosphatase activity between 4 and 6 hours, an increase that has been shown to be strongly correlated with formation of the pre-spore septum during stage II of sporulation (Errington, 2003; Glenn & Coote, 1975). Schujman et al. (1998) hypothesized that the mother cell-specific transcription factor, σ^E^, might be formed by proteolytic cleavage of pro-σ^E^ only in that compartment via a mechanism dependent on the asymmetric composition of pre-spore septum phospholipids. Moreover, Ruzal et al. (1998) showed that hyperosmotic media blocked sporulation at the σ^E^ activation stage, while López et al. (1998) found that *B. subtilis* cells grown in hyperosmotic medium exhibited a marked decrease in iso-branched fatty acids. Thus, it is logical to hypothesize that the mother-cell side of the forespore membrane is enriched in iso-branched fatty acids, especially 13-methyltetradecanoic acid (iC15). This simple mechanism appears to have the potential to finally resolve the question as to how this asymmetric septum functions as “an organelle for the establishment of cell fate” (Losick & Dworkin, 1999).

## Experimental Procedures

### Bacterial strains and growth conditions

Most of the work was done with the prototypical *Bacillus subtilis ΔtrpC2* auxotroph (168 strain), obtained from the National BioResource Project (NBRP; Shizuoka, Japan; Strain No. MGNA-A001). Three mutant *B. subtilis* strains obtained from the *Bacillus* Genetic Stock Center (Columbus, OH, USA) were also studied, namely: *Δspo0A::ermtrpC2* (BKE24220), *ΔfadN::ermtrpC2* (BKE32840) and *ΔmmgA::ermtrpC2* (BKE24170). Bacteria were maintained as glycerol stocks at-80°C.

Inoculum cultures were grown overnight in 25 g L^-1^ Lysogeny Broth powder plus 5.0 g L^-1^ dextrose, with shaking at 250 rpm at 37 °C. Non-sporulation medium of 100 ml volume in a one-liter flask had the same composition but was buffered at an initial pH of 7.0 by supplementing with 2.30 g L^-1^ NaH_2_PO_4_·H_2_O and g L^-1^ Na_2_HPO_4_·7H_2_O. It was inoculated at 10 %v/v and incubated as above. Sporulation medium of 100 ml volume was as per Leighton and Doi (1971), consisting of salts plus Bacto Peptone and Beef Extract Powder as carbon and nitrogen sources, with only 1.0 g L^-1^ of dextrose, inoculated at 5 %v/v and incubated as above.

Culture medium was supplemented with one or more of the following, as stated in the text: 50 mg of *L*-valine-d_8_, or an equimolar amount of *L*-valine or *L*-valine-^13^C_5_,^15^N; 25 mg of hexanoic-d_11_ acid; or 50 mg of *D,L*-methionine-3,3,4,4-d_4_ or *D,L*-methionine. *L*-valine-d_8_, hexanoic-d_11_ acid and *D,L*-methionine-3,3,4,4-d_4_ were obtained from C/D/N Isotopes Inc., Pointe Claire, QC, Canada; *L*-valine-^13^C_5_,^15^N was from Cambridge Isotope Laboratories, Inc., Tewksbury, MA, USA.

One culture was performed with *Xanthomonas campestris* ATCC 33440, an inoculum culture of which was grown for three days with shaking at 200 rpm and 30 °C in yeast extract-casamino acids medium (Blanvillain et al., 2007). Fifty ml of the same medium, supplemented with 0.5 mg ml^-1^ of 2-methylpropionic-d_7_ acid (C/D/N Isotopes Inc.), was neutralized, inoculated at 5 %v/v and incubated as above. After 48 hours the biomass was harvested and prepared for analysis of cellular fatty acids as described below.

### Biomass harvest and extraction

For cultures on non-sporulation medium destined for acyl-CoA analysis, biomass was harvested 1.5-2.0 h after the onset of stationary phase as indicated by minimum broth pH (Hanson et al., 1963; i.e., following 6-7 h of culture) by vacuum filtration through 0.45 µm nylon membranes, then transferred into 5.0 ml of ice-cold acyl-CoA extraction solvent (Bennett et al., 2009), containing 4.0 μg ml^-1^ of benzoyl-CoA as internal standard. The suspension was alternately vortexed and immersed in ice for 5 min, then stored at-80 °C.

For cultures on sporulation medium destined for fatty acids or acyl-CoA analysis, aliquots of broth were harvested every hour from 3 to 6 hours of culture, equally divided between two tubes and centrifuged for 10 min at 17,000 *g* at 4 °C. The clear supernatant was pooled, and an aliquot stored at-20 °C for subsequent IRMS analysis (see below). The bacterial pellet was washed by centrifugation with 10 ml of 18 Milli-Q water, and the supernatant was poured off then quantitatively removed by pipet. Each pellet was suspended in 400 μl of absolute ethanol containing 20 μl of concentrated sulfuric acid and 20 μg ml^-1^ of *n*-tridecanoic acid as internal standard. In some cases, one pellet was suspended in acyl-CoA extraction solvent containing benzoyl-CoA as above. The suspensions were stored at-80 °C until further processed. Tubes containing the suspension in acyl-CoA extraction solvent were centrifuged for 5 min at 16,000 *g*, then the clear supernatant was transferred to a new tube and its volume reduced via SpeedVac with heat for 30 min. Its volume was made to 600 μl with Milli-Q water before filtration through a 0.2 μm nylon membrane for HPLC analysis.

For culture samples destined for biomass amino acid analysis, approximately 25 ml of broth was divided between two 15-ml tubes and centrifuged for 10 min at 3,900 *g* at 4 °C. The supernatant was poured off, each pellet rinsed by vortexing with 500 μl of 0.82 % NaCl solution, the suspension combined in a new 1.5 ml conical polypropylene tube and centrifuged at 18,800 *g* for 5 min at 4°C. The supernatant was removed by aspiration and the pellet stored at-20°C, pending derivatization and amino acid analysis.

### Fatty acid methyl ester preparation and GC-MS analysis

Fatty acid standards myristic (*n*-tetradecanoic) acid and isopalmitic (14-methylpentanoic) acid were obtained from Sigma-Aldrich, Oakville, ON, Canada; *n*-tridecanoic acid was from Acros Organics, Geel, Belgium; hexane and petroleum ether (80-110 °C boiling range) were analytical grade.

Biomass fatty acid methyl esters were prepared and analyzed using procedures adapted from Long and Antoniewicz (2014). The acidic methanol biomass suspension was hydrolysed at 100 °C, then extracted into 750 μl of hexane or petroleum ether. The solvent was evaporated via SpeedVac for 15 min with heat, the residue dissolved in 150 μl or 160 μl of the same solvent, and the solution placed in a 250 μl glass insert vial. GC/MS analysis was performed on a Trace GC Ultra gas chromatograph with AS3000 Autosampler equipped with a Zorborbax DB-5 column (30 m, 0.25 mm i.d., 0.20 μm film thickness; Agilent J&W Scientific), with the eluate transferred at 280 °C to a Polaris Q mass spectrometer (Thermo Scientific). GC-MS conditions were as per Long and Antoniewicz (2014), except that ion source temperature was 250 °C, helium flow rate was 1.1 ml min^-1^, and the following shortened temperature program was generally used: initial temperature of 80 °C, increased to 160 °C at 50 C° min^-1^ then to 230 °C at 10 C° min^-1^, finally to 310 °C at 50 C° min^-1^, and held for 3 min.

For each sample and each scanning mode, six injections were performed, intercalated with the other samples from the same culture. Initial work employed only a scan of *m/z* values from 60-360 from 4-8 min in positive-ion mode. To obtain more accurate mass isotopomer data, later work also analyzed sets of injections using two distinct segmented-scan methods, as follows:

a. Scan *m/z* 227-233 [i.e. (M-1) to (M+5)] from 4.5 min for the nC13 internal standard eluting at 4.64 min; *m/z* 248-254 from 4.8 min for iC14-d7 eluting at 5.10 min; and *m/z* 276-282 from 6.0 min for iC16-d7 eluting at 6.82 min.
b. Scan *m/z* 227-233 from 4.6 min for nC13; *m/z* 241-247 from 5.0 min for iC14 and nC14 eluting at 5.15 min and 5.45 min, respectively; *m/z* 255-261 from 5.7 min for iC15 and aC15 eluting at 6.0 and 6.08 min, respectively; *m/z* 269-275 from 6.5 min for iC16 and nC16 eluting at 6.87 min and 7.2 min, respectively; and *m/z* 283-289 from 7.4 min for iC17 and aC17 eluting at 7.75 and 7.85 min, respectively.

### HPLC/MS analysis of acyl-CoA esters

CoA ester analysis was performed in MRM mode on a Waters Model 2795 Separation Module connected to a Waters Quattro Premier XE MicroMass mass spectrometer running on MassLynx V4.1 software. Separation was performed on a Gemini C_18_ analytical column (150 x 3 mm, 5 µm particle size, 110 Å pore size; Phenomenex) equipped with a C_18_ SecurityGuard column, using 10 mM ammonium acetate pH 5.0 in either water (eluent A) or acetonitrile (eluent B) at a flow rate of 400 µl min^-1^. The following optimized gradient program was used (% of eluent A): 96% for 1 min, linearly to 60% at 7 min, to 50% at 9 min, to 0% at 11 min and holding until 15 min; then to 96% at 16 min and holding until 20 min. At least six 50 µl injections were done for each sample.

Approximately one-tenth of the column eluent was fed into the Z-spray ionization interface of the mass spectrometer operated in positive-ion mode, using collision energy of 30 V and cone voltage of 50 V. The flow rate of the ultra-high purity argon collision gas was 0.30 ml min^-1^ at a pressure of 3.7 x 10^-3^ mbar. The *m/z* values of the precursor and major daughter (quantifier) ions for each acyl-CoA, determined from the mass spectrum of the pure standards, are shown in Table 3.

**TABLE 3.** LC/MS/MS parameters of selected acyl-CoA esters.

| Acyl-CoA ester | Retention time, min | Precursor ion $m/z$ | Quantifier ion(s) $m/z$ |
| --- | --- | --- | --- |
| Propionyl-CoA | 6.7 | 824.1 | 214.9, 317.9 |
| Propionyl-d<sub>5</sub>-CoA | 6.7 | 829.1 | 322.9 |
| Isobutyryl-CoA | 7.3 | 838.1 | 228.9, 331.9 |
| Isobutyryl-d<sub>7</sub>-CoA | 7.3 | 845.1 | 235.9, 338.9 |
| 2-Methylbutyryl-CoA | 8.0 | 852.1 | 242.9, 345.6 |
| 2-Methylbutyryl-d<sub>9</sub>-CoA | 8.0 | 861.1 | 251.9, 354.6 |
| Benzoyl-CoA | 8.2 | 872.1 | 365.2 |

### GC/MS analysis of amino acids

Amino acids were analyzed using a method adapted from Dauner and Sauer (2000). The biomass pellet was acid hydrolyzed at 105 °C, then evaporated to dryness by adding 500 µl of anhydrous ethanol to each of two tubes and heating on a SpeedVac under vacuum for 30 min. Aliquots of ethanol were added and evaporation continued until a total of ≥ 2 ml had been added to each tube, and the solids were apparently dry. Finally, the samples were made anhydrous by addition of 50 µl of dichloromethane and SpeedVac evaporation for 10 min. The contents of one tube were dissolved in 80 or 100 μl of THF and derivatized with an equal volume of MTBSTFA, then transferred to a glass insert vial for analysis.

GC/MS analysis was performed as described above, except that the temperature program used was 150 °C for 2 min, 3 C°/min to 240 °C, and 10 C°/min to 320 °C, scanning from *m/z* 150-490. The valine peak eluted from 7.85-8.02 min, containing mother-ion fragments of *m/z* 186, 260 and 288. The fraction of carbon-13 labeled valine was calculated based on 12 injections by dividing the sum of the *m/z* 190, 191, 264, 265 and 293 intensities by the sum of those from *m/z* 186-191, 260-265 and 288-293.

### Miscellaneous methods

Culture broth optical density at 600 nm was converted to bacterial dry mass concentration using the estimate of 0.39 g L^-1^ for *E. coli* from a standard database (http://bionumbers.hms.harvard.edu; ID#109836), since no corresponding datum for *B. subtilis* is available. Thus, the average absorbance at 600 nm was multiplied by 0.39 to estimate biomass concentration in g dry matter L^-1^.

Isotope ratio mass spectrometry (IRMS) for measurement of the deuterium oxide content of culture broth water was performed by the Light Stable Isotope Geochemistry Laboratory, Research Centre on the Dynamics of the Earth System, University of Quebec at Montreal (Geotop-UQAM, Montreal, QC, Canada). Exactly 2.00 ml of sample water was pipetted into a 3 ml vial, closed with a septum cap and transferred to a 40°C heated rack. A hydrophobic platinum catalyst (Hokko beads) was added. After 1 hour, air in the vials was replaced with H_2_ using the AquaPrep. Samples were left to equilibrate for 4 hours. The equilibrated samples were analyzed with a Micromass model Isoprime isotope ratio mass spectrometer coupled to an AquaPrep system in dual inlet mode. Three internal reference waters (δ^2^H = 1.28 ± 0.27‰; 98.89 ± 1.12‰;-155.66 ± 0.69‰) were used to normalize the results on the VSMOW-SLAP scale. A 4^th^ reference water (δ^2^H =-25.19 ± 0.83‰) was analyzed as an unknown to assess the exactness of the normalization. Results were given in delta units (δ) in ‰ vs VSMOW. The overall analytical uncertainty is better than ±2.0 ‰ for δ^2^H. This uncertainty is based on the propagation of uncertainties of the normalization of the internal reference materials and the samples.

### Calculation of acyl-CoA ester concentrations

HPLC/MS analysis of acyl-CoA esters was calibrated by determining the response ratio of the analyte to the benzoyl-CoA internal standard over a suitable range of concentrations of the former. Specifically, 6 injections were done at each of three analyte concentrations with fixed benzoyl-CoA concentration of 10 μg ml^-1^, and the response ratio calculated for each as follows: (total area of analyte quantifier fragment peaks) x (benzoyl-CoA concentration) / (area of benzoyl-CoA fragment). The mean and standard deviation of this ratio were calculated for each analyte concentration, and any outliers removed using Pierce’s criterion (Ross, 2003). The resulting mean ratios were plotted against analyte concentration to derive the response ratio slope (giving correlation coefficients of 0.997-0.999).

To calculate sample acyl-CoA concentrations, the peak area for the quantifier ion(s) of each analyte and the internal standard were tabulated for each injection; when two quantifier ions were used, their sum was calculated. An internal standard ratio for each injection was calculated as the expected benzoyl-CoA concentration divided by the area of its quantifier peak. Analyte concentration in the solution injected was then calculated, for each injection, as follows: (internal standard ratio) x (total area of analyte quantifier fragment peaks) / (analyte response ratio slope), μg ml^-1^. The mean and standard deviation of this value were calculated, and any outliers removed as described above. The resulting mean and standard deviation were converted to molar concentration, and finally the cytoplasmic concentration was estimated based on the bacterial dry matter (DM) present in the broth sample extracted, and assuming of an intracellular liquid phase volume of 1.5 µl mg^-1^ DM.

### Calculation of fatty acid concentrations

Using data from the *m/*z 60-360 scan injections, the total ion count for each fatty acid and internal standard was tabulated for each injection. The ratio of its count to that of the internal standard was calculated for each injection, along with the mean ratio and its standard deviation. Next, Pierce’s criterion (Ross, 2003) was used to detect and remove any outliers, resulting in at least 4-6 values for recalculation of each mean ratio and standard deviation. This was converted to the mean concentration and standard deviation of each fatty acid in the analytical solution (μg μl^-1^) based on the internal standard mass nominally present. These values were converted to μg mg^-1^ bacterial dry matter (DM) based on the mass of the latter estimated to have been present in the broth sample as calculated from its optical density at 600 nm. Finally, the standard deviation of the mean concentration was calculated based on the actual number of ratio values used to calculate the mean.

Using data from the two segmented-scan protocols, the sum of the intensity of the (M-1) to (M+5) peaks was calculated for each injection of each fatty acid, and any outliers removed as described above. The resulting mean intensity for each fatty acid was summed to give the total intensity of the ten fatty acids analyzed, and each as a percentage of the total calculated. These values for each fatty acid at each sampling time were finally averaged over the triplicate cultures for each treatment studied, and the standard error calculated.

### Mass isotopomer distribution analysis

#### Natural abundance mass isotopomer (analog) distributions

The theoretical mass isotopomer distribution (MID) vector for each fatty acid was calculated as described by González-Antuña et al. (2014) via their IDC.xls spreadsheet using a value of 1.09% carbon-13 natural abundance. The elements of this vector, denoted *A^i^*, were calculated as the ratio of the peak intensity of the respective isotopomer to the sum of the M (lowest molecular weight “mother” ion) to (M+9) peak intensities, with ∑ A^i^ = 1. The theoretical natural abundance MID for iC14-d7 and iC16-d7 fatty acids was calculated similarly using a revised spreadsheet explicitly incorporating deuterium as an element.

#### MID concentration dependence analysis

Given an observed concentration dependence of fatty acid methyl ester isotope ratios using electron impact ionization on some mass spectrometers (Fagerquist et al., 1999; González-Antuña et al., 2014; Patterson and Wolfe, 1993), we analyzed and attempted to correct our raw data for such an effect. Specifically, using 3h data for each fatty acid from triplicate cultures of the *ΔtrpC2* strain supplemented with *L*-valine, the intensities of the (M+1) and (M+2) analogs were plotted versus the intensity of M and fitted to a second-order regression equation with zero intercept (Patterson and Wolfe, 1993). Using the parameters so derived, the apparently “excess” (M+1) and (M+2) intensities were subtracted from the observed values by extrapolation to zero fatty acid concentration (ibid.). The theoretical natural abundance intensity for both analogs was also calculated based on the observed M intensity and the theoretical MID of each fatty acid. The latter values were greater than those derived from the “excess” subtraction procedure for 99 per cent of the injections for every fatty acid except aC15, whose anomalous MID results obtained via the *m/*z 60-360 scan protocol were excluded from consideration. Hence, we concluded that the raw data did not require adjustment for concentration dependence.

#### Calculation of cluster purity parameters

The parameters for cluster purity, *X_M-1(H)_* and *X_M_*, arising from loss of hydrogen from the mother (M) ion and/or tailing of M into the (M-1) peak in the experimental spectrum, were calculated by solution of Equation (1) using multiple linear regression. Each *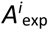* is the average experimentally observed element of the MID vector after removal of any outliers using Pierce’s criterion (Ross, 2003), with ∑ A^i^ = 1. Similarly, each 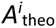 is the corresponding element of the theoretical mass isotopomer distribution vector, calculated as described above.

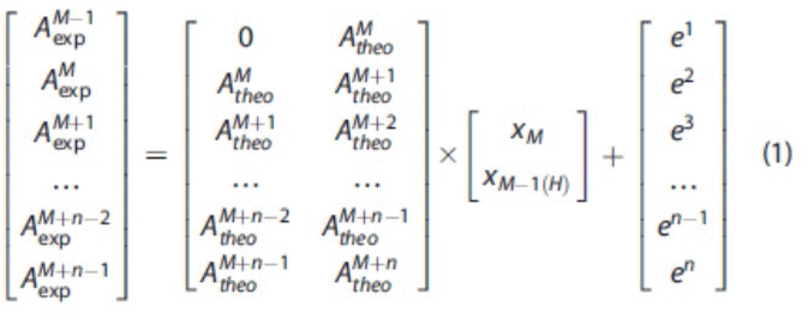

In practice, we augmented the “Brombuterol LC.xls” spreadsheet of González-Antuña et al. (2014) to accept data from six injections rather than five, and processed the 3h data for each fatty acid from triplicate cultures of the *ΔtrpC2* strain supplemented with *L*-valine. Thus, using the corresponding theoretical MID, we calculated mean values of *X_M-1(H)_* and *X_M_*, and a standard error of the mean, *Ux_M-1(H)_* for each fatty acid.

A similar but slightly revised procedure was used to calculate cluster purity parameters for iC14-d7 and iC16-d7 fatty acids, using 3h data from 10 cultures of various *B. subtilis* strains supplemented with *L*-valine-d_8_ (i.e. *ΔtrpC2*, *Δspo0A::ermtrpC2*, *ΔfadN::ermtrpC2*, and *ΔmmgA::ermtrpC2*). This procedure used only the (M_d7_-1) to (M_d7_+3) data to avoid interference in the calculation by the high intensity of the (M_d7_+4) analog.

#### Calculation of convoluted mass isotopomer distributions

To correct the experimentally observed mass isotopomer distributions for cluster purity and lack of spectral resolution we calculated so-called “convoluted” analog distributions as described by González-Antuña et al. (2014). First, for each segmented-scan injection and each fatty acid we calculated 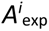 as the ratio of the peak intensity of each mass divided by the sum of the measured (M-1) to (M+5) peak intensities (in early work, (M-1) to (M+9)). Any outliers were removed using Pierce’s criterion (Ross, 2003), and the mean and standard deviation calculated. Then, using the cluster purity parameters *X_M-1(H)_* and *X_M_* calculated for each fatty acid, we calculated a convoluted isotope distribution, 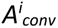, according to Equation (2), and the standard deviation for each analog by statistically combining the uncertainty *Ux_M-1(H)_* with that of the mean 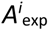 value.

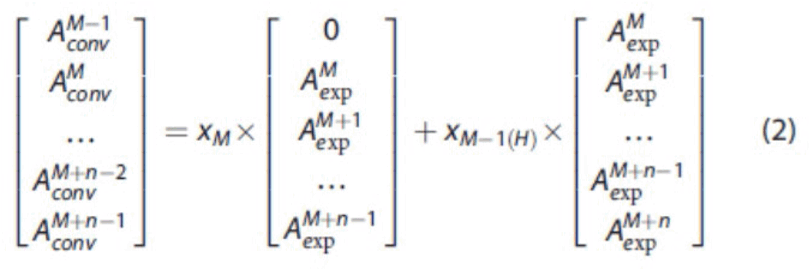

Finally, the values of 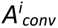 for each fatty acid were summed and the values in the distribution normalized to total exactly one to give the uncorrected distribution vector, *MIDV_un_*; the standard deviations were normalized identically, as shown in the appended files “RC129 iC14,16-d7 distribution calculations.xlsx” and “RC129 other FA distribution calculations.xlsx”.

#### Calculation of mass isotopomer enrichment

For each “treatment” (i.e., different labeled substrate added), and for each sampling time (i.e., 3, 4, 5 and 6 h), the values in the normalized analog distributions (*MIDV_un_*) from triplicate cultures were averaged and the standard error of the mean, SEM_MIDVun_, calculated. These mean distributions were then corrected for natural abundance via the correction matrix method (Midani et al., 2017; Nanchen et al., 2007) as described in detail below, and as shown in file “Valine-d8 enrichment calculations.xlsx”.

A natural abundance correction matrix (*CM*) for each fatty acid, including iC14-d7 and iC16-d7 fatty acids, was created from the theoretical mass isotopomer distribution vector, calculated as described above. The inverse of each correction matrix (*CM*^-1^) was calculated and multiplied by the normalized distribution vector, *MIDV_un_*, to give *MIDV_c_*. Any (very occasional) negative value of the latter was set to zero, and the elements of the vector were summed and normalized as above to give the final corrected distribution vector, *MIDV_c_*, comprised of elements designated *A_c_ ^i^*. The fractional labeling (% enrichment), if any, of each mass isotopomer of each fatty acid was then calculated according to Equation (3), where *i* is the shift of isotopomer mass relative to monoisotopic mass (M), and *N* is the number of carbon atoms in the fatty acid without the methoxy group added by esterification (Nanchen et al., 2007; Wu et al., 2020).

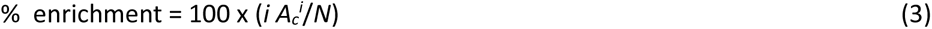

The uncertainty in the fractional labeling values was calculated according to the method given by Von Freese (2020), specifically starting with Equation (4), where σ represents standard deviation and *CM*^t^ is the transpose of a given *CM*.

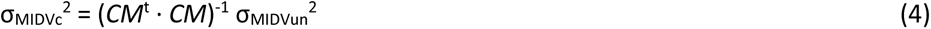

Finally, the square root of σ_MIDVc_^2^ was taken, and the standard error of each % enrichment value calculated using Equation (3) with σ(*A_c_ ^i^*) in place of *A_c_ ^i^*. Only enrichment values which exceeded zero by at least three times the standard error were considered sufficiently significant to include in the data reported in the Results section.

## Acknowledgments

The authors wish to acknowledge the generous assistance of Sylvain Milot with mass spectrometric methodology, Marie-Christine Groleau for microbiological assistance, and Arvin Nickzad for help in compiling some of the GC-MS results.

## Appendix 1

**Table A1:** Percentage of various fatty acids versus time for replicate B. subtilis cultures of various strains analyzed using m/z 60-360 scan mass spectrometry^a^.

|  | 3h |  | 4h |  | 5h |  | 6h |  |
| --- | --- | --- | --- | --- | --- | --- | --- | --- |
| Fatty acid | Mean | SEM | Mean | SEM | Mean | SEM | Mean | SEM |
| iC15 | 11.8 | 0.3 | 14.2 | 0.6 | 19.3 | 0.8 | 23.2 | 0.6 |
| Total iso | 34.4 | 1.1 | 36.9 | 1.1 | 42.0 | 1.2 | 46.4 | 1.2 |
| Total anteiso | 52.4 | 1.5 | 49.5 | 1.6 | 48.0 | 1.3 | 45.6 | 1.2 |
| Total normal | 11.3 | 0.8 | 10.1 | 0.7 | 8.4 | 0.5 | 7.1 | 0.4 |

<sup>a</sup> Strains: $\Delta trpC2$ , $\Delta spo0A::ermtrpC2$ and $\Delta fadN::ermtrpC2$ ; n=19.

**Table A2:** Cellular fatty acids concentration (microgram g^-1^ DM) versus time for replicate *B. subtilis ΔtrpC2* cultures supplemented with labeled or unlabeled *L*-valine; quantified using *m/z* 60-360 scan mass spectrometry.

|  | 3h |  | 4h |  | 5h |  | 6h |  |
| --- | --- | --- | --- | --- | --- | --- | --- | --- |
| Fatty acid | Mean | SEM<sup>a</sup> | Mean | SEM<sup>a</sup> | Mean | SEM<sup>a</sup> | Mean | SEM<sup>a</sup> |
| iC14-d7 | 5 | 2 | 5 | 1 | 11 | 1 | 13 | 2 |
| iC14 | 13 | 2 | 11 | 2 | 11 | 2 | 10 | 2 |
| nC14 | 11 | 2 | 9 | 1 | 4 | 1 | 6 | 1 |
| iC15 | 274 | 20 | 279 | 24 | 366 | 24 | 425 | 33 |
| aC15 | 695 | 42 | 559 | 50 | 523 | 36 | 523 | 38 |
| iC16-d7 | 23 | 7 | 19 | 0 | 36 | 5 | 39 | 5 |
| iC16 | 176 | 26 | 135 | 26 | 130 | 24 | 141 | 25 |
| nC16 | 249 | 23 | 176 | 23 | 137 | 21 | 124 | 15 |
| iC17 | 352 | 35 | 280 | 44 | 256 | 31 | 273 | 28 |
| aC17 | 418 | 43 | 310 | 50 | 242 | 29 | 238 | 23 |
| Total | 2217 | 202 | 1782 | 221 | 1716 | 174 | 1794 | 172 |

<sup>a</sup> n=3 for iC14-d7 and iC16-d7; otherwise, n=9.

**Table A3:** Percentage 3-hour enrichment of replicate B. subtilis ΔtrpC2 and Δspo0A::ermtrpC2 cultures supplemented with L-valine-d_8_.

|  |  | *$\Delta trpC2$* |  | *$\Delta spo0A::ermtrpC2$* |  |
| --- | --- | --- | --- | --- | --- |
|  | Analog | Mean | SEM<sup>a</sup> | Mean | SEM<sup>b</sup> |
| **iC14-d7** | M+5 | 2.25 | 0.05 | 2.32 | 0.02 |
|  | M+4 | 8.56 | 0.14 | 9.97 | 0.17 |
|  | M+3 | 0.00 | 0.06 | 0.00 | 0.07 |
|  | M+2 | 0.63 | 0.13 | 0.60 | 0.09 |
|  | M+1 | 0.54 | 0.05 | 0.45 | 0.13 |
| **iC16-d7** | M+5 | 1.03 | 0.04 | 0.48 | 0.01 |
|  | M+4 | 2.86 | 0.11 | 1.46 | 0.07 |
|  | M+3 | 0.00 | 0.04 | 0.00 | 0.01 |
|  | M+2 | 0.21 | 0.02 | 0.22 | 0.02 |
|  | M+1 | 0.93 | 0.09 | 0.99 | 0.11 |

<sup>a</sup> n=3; <sup>b</sup> n=2.

**Figure A1(a):**
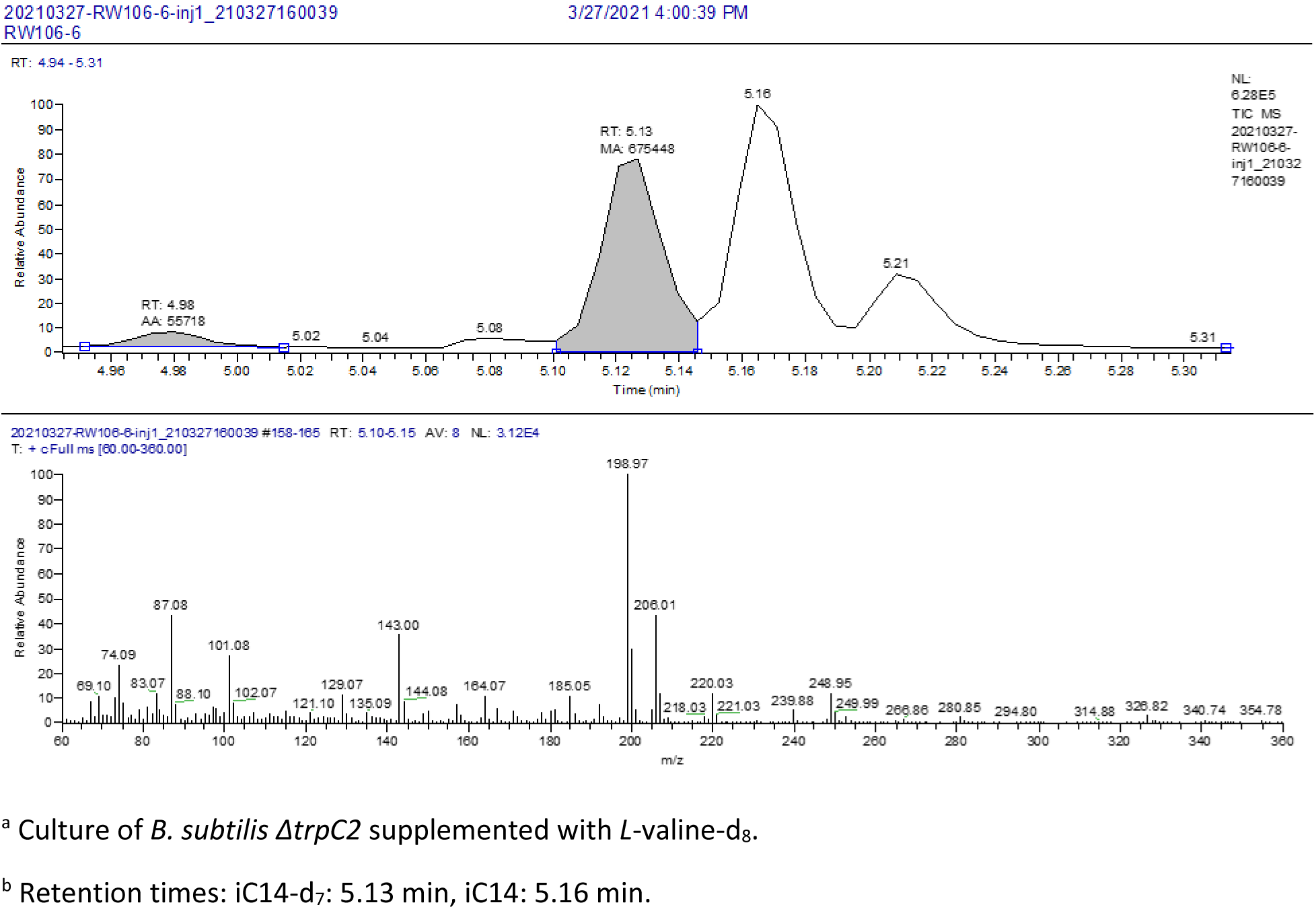
Chromatogram showing separation of *B. subtilis* iC14-d_7_ (shaded, “RT: 5.13”) from iC14 (“5.16”), and the iC14-d_7_ mass spectrum^a,b^.

**Figure A1(b):**
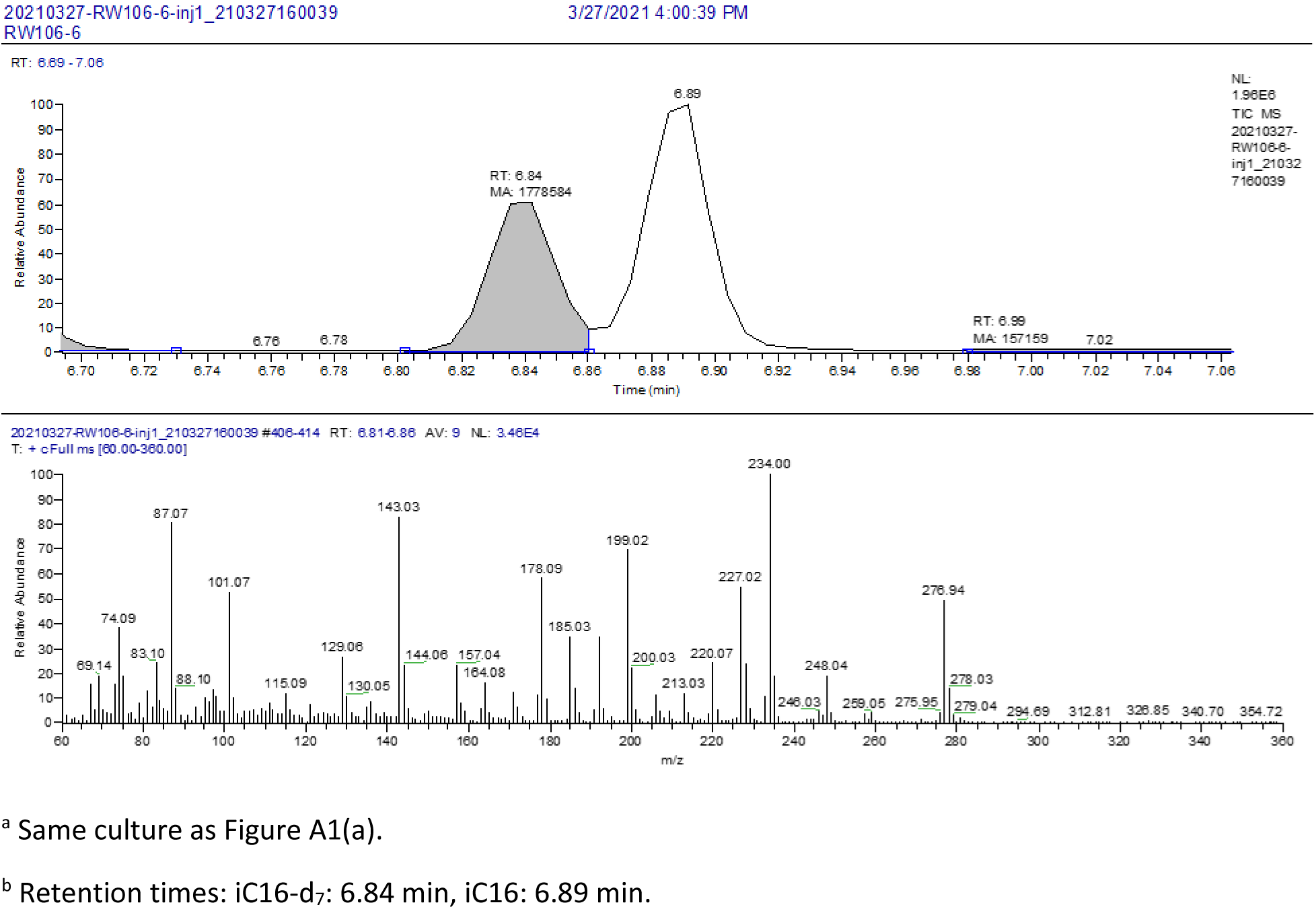
Chromatogram showing separation of *B. subtilis* iC16-d_7_ (shaded, “RT: 6.84”) from iC16 (“6.89”), and the iC16-d_7_ mass spectrum^a,b^.

**Figure A1(c):**
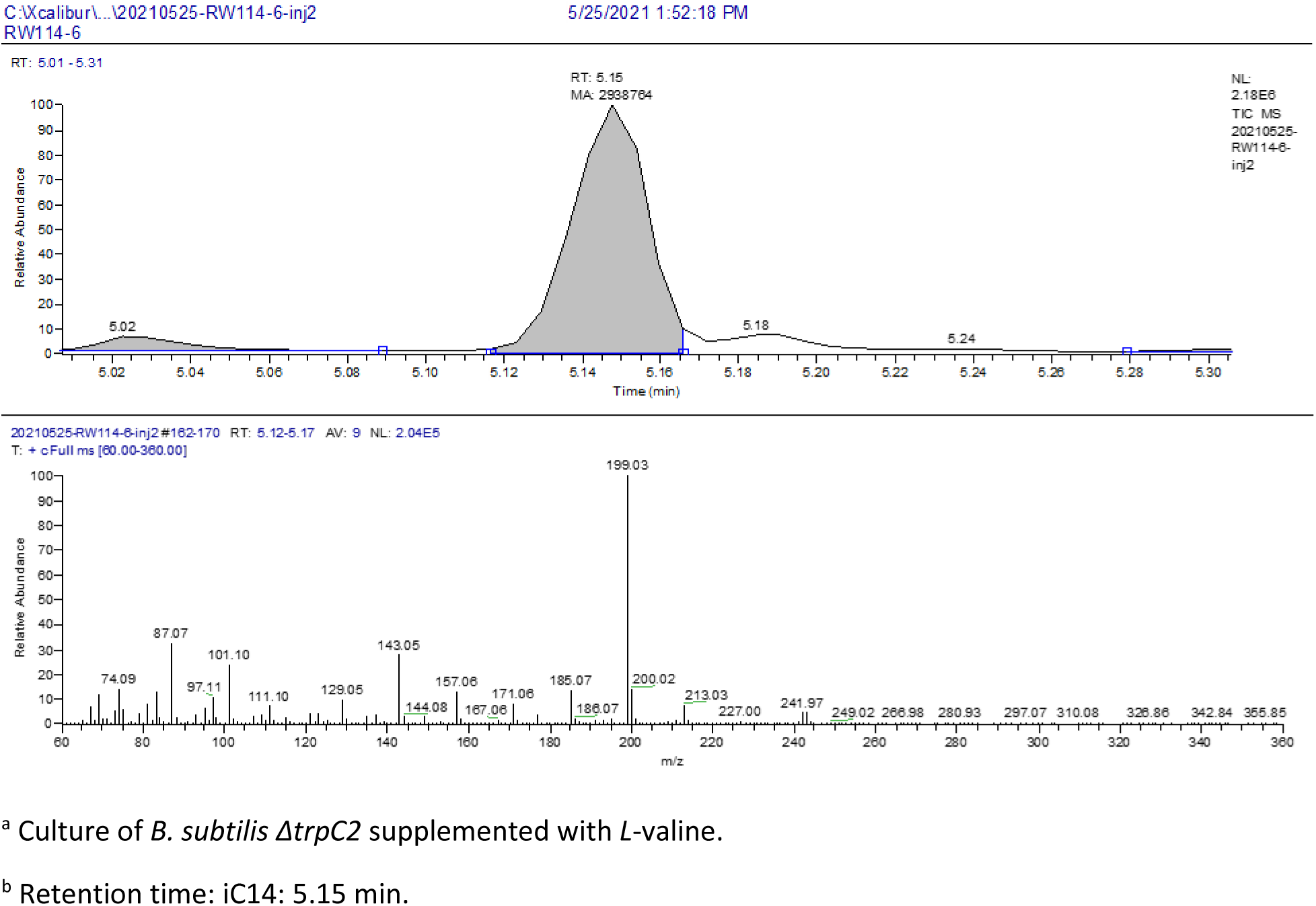
Chromatogram of *B. subtilis* iC14 (shaded, “RT: 5.15”), and the 5.12-5.17 minute iC14 mass spectrum^a,b^.

**Figure A1(d):**
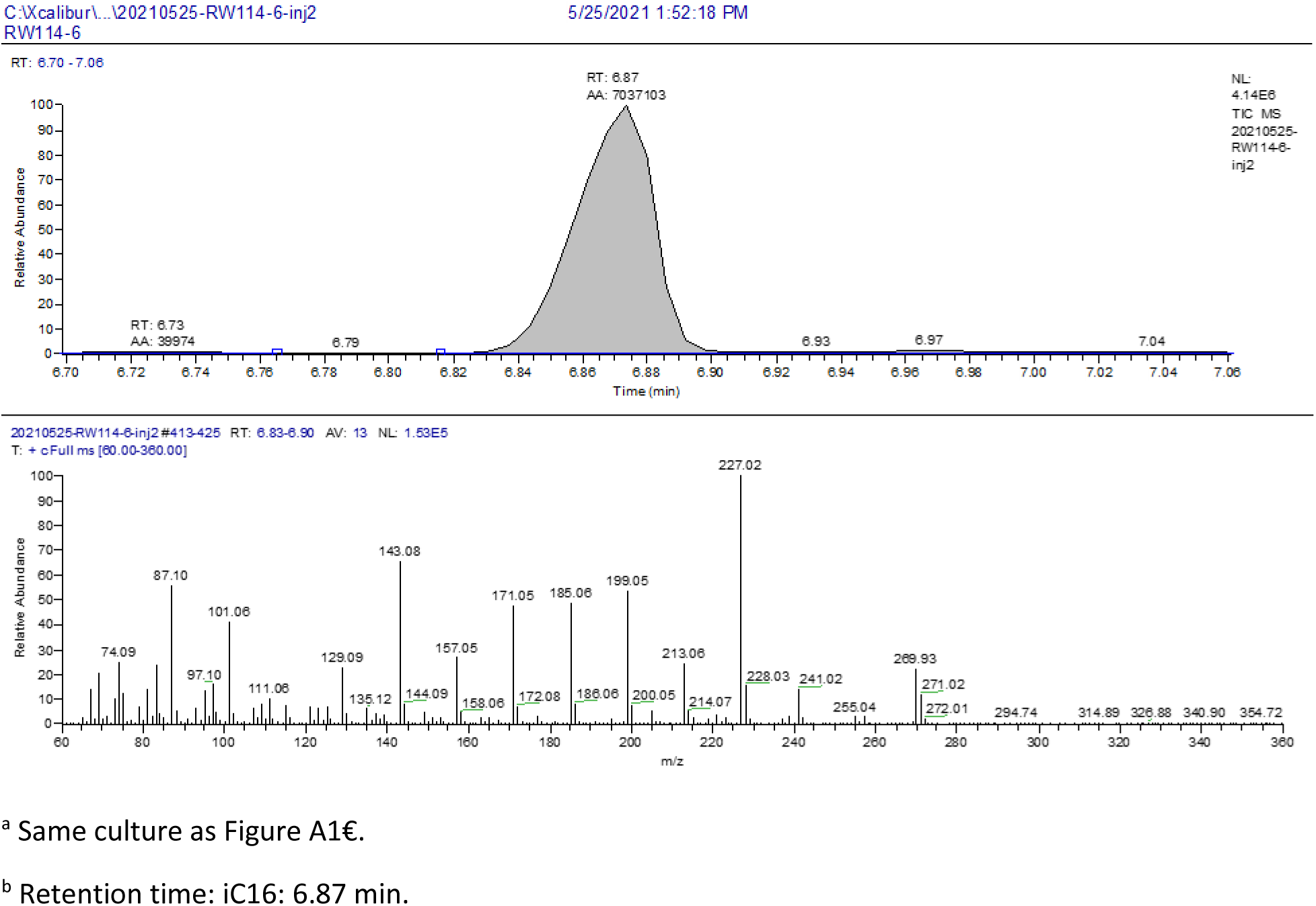
Chromatogram of *B. subtilis* iC16 (shaded, “RT: 6.87”), and the 6.83-6.90 minute iC16 mass spectrum^a,b^.

**Figure A1(e):**
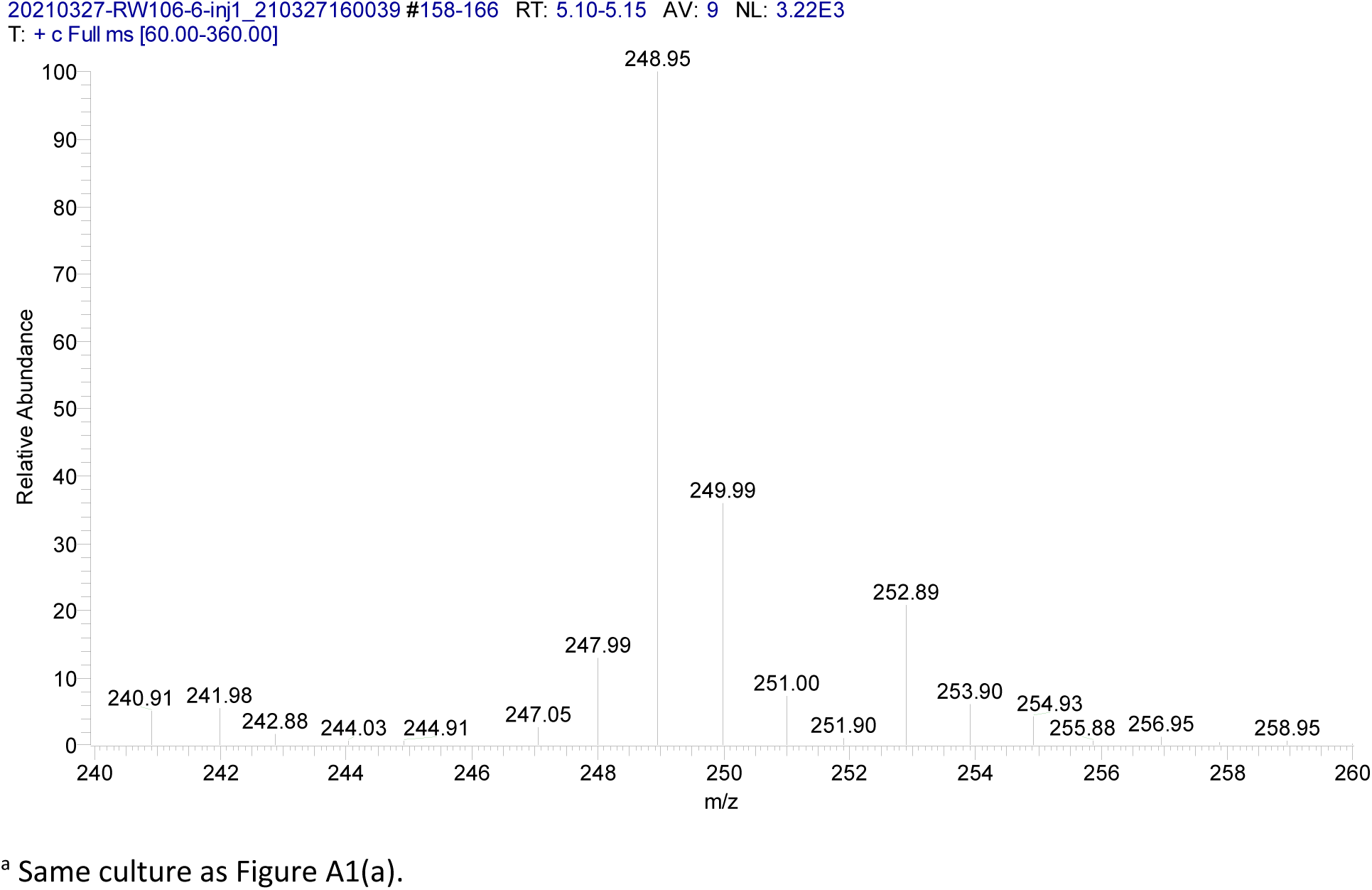
Detail of the mass spectrum of *B. subtilis* iC14-d ^a^.

**Figure A1(f):**
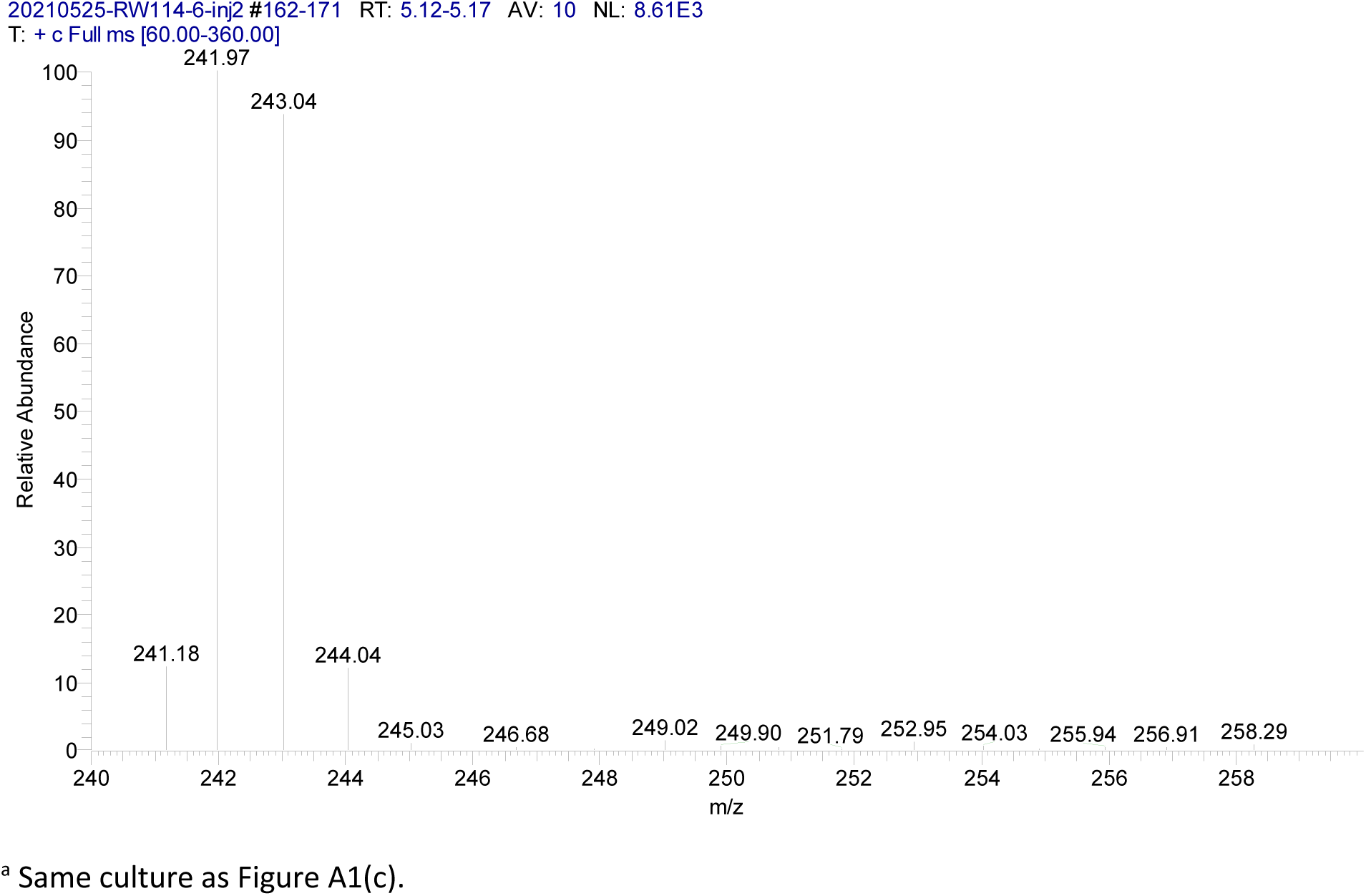
Detail of the mass spectrum of *B. subtilis* iC14^a^.

**Figure A2(a):**
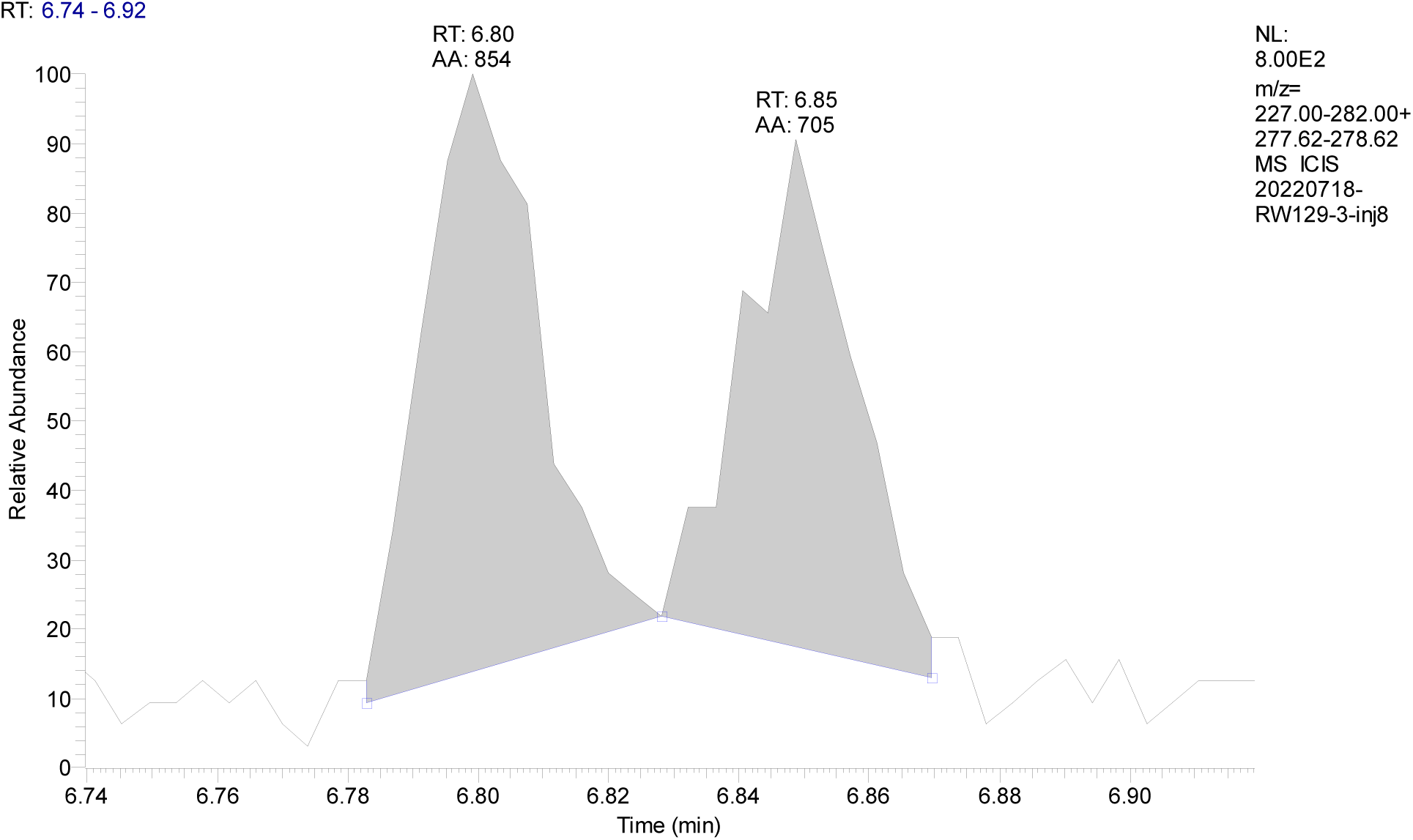
Chromatogram of a typical composite iC16-d7 fatty acid peak in a *B. subtilis ΔtrpC2* culture supplemented with *L*-valine-d_8_, scanned from *m/z* 276 to 282.

**Figure A2(b):**
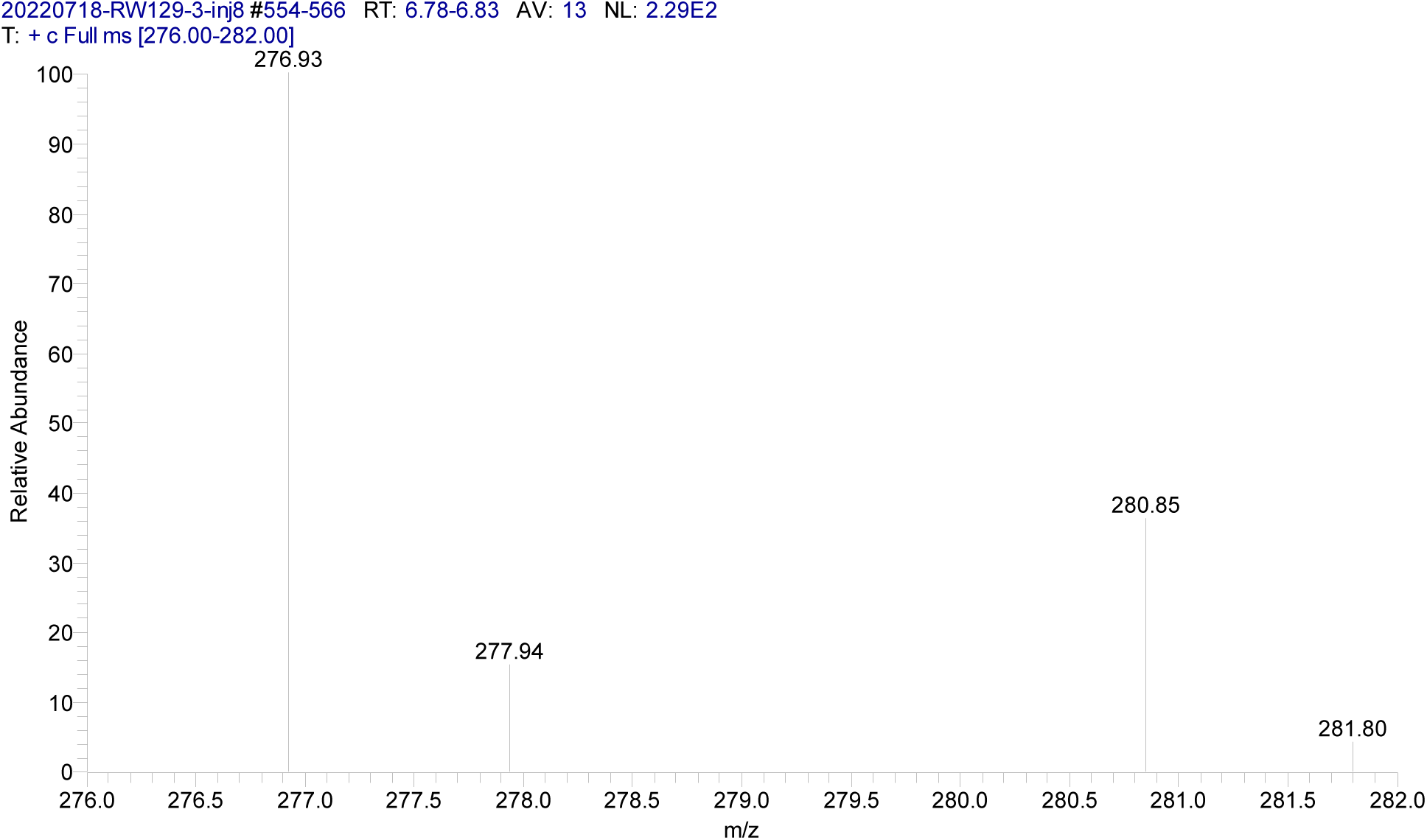
Mass spectrum of the leading *B. subtilis* 6.78-6.83 minute peak.

**Figure A2(c):**
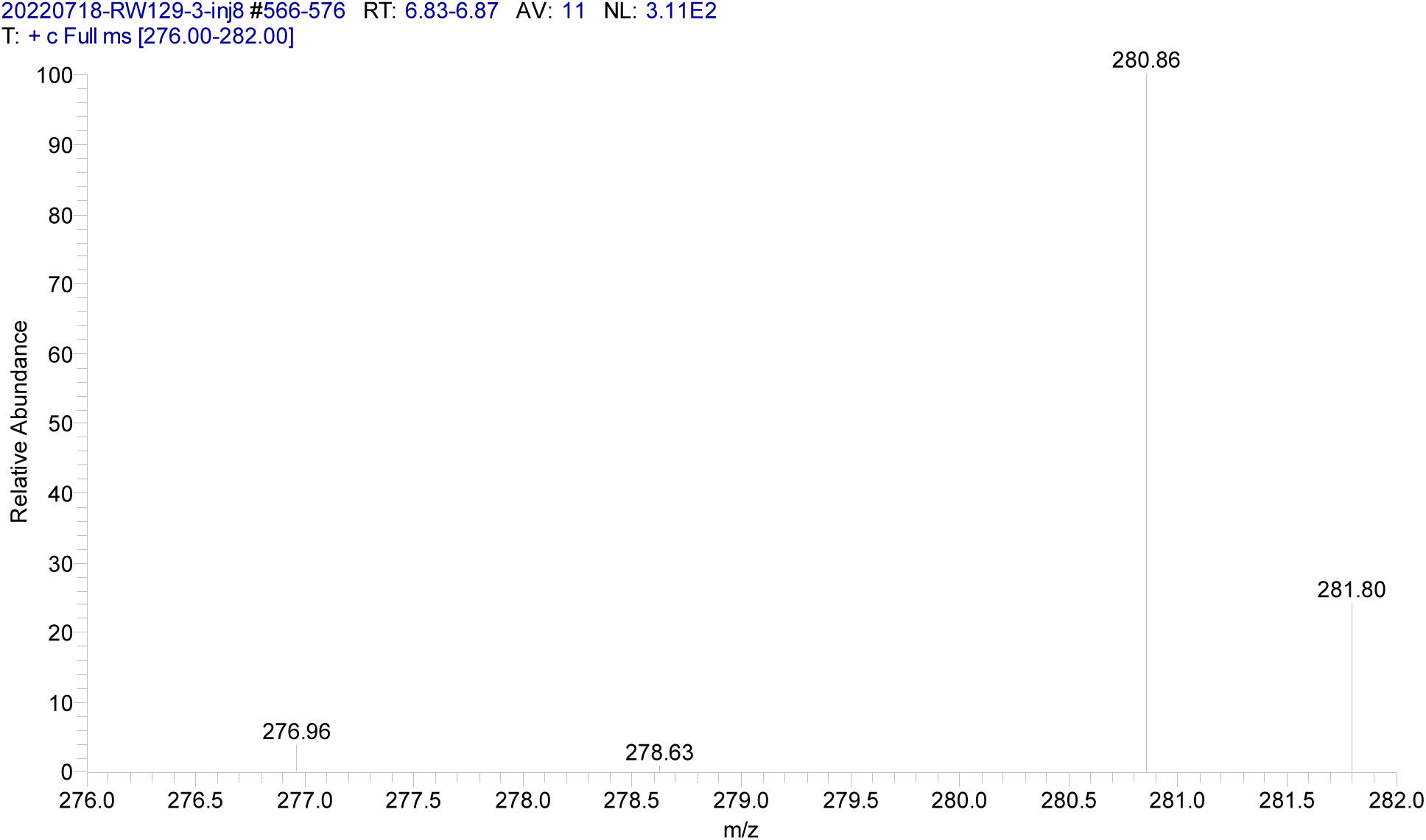
Mass spectrum of the trailing *B. subtilis* 6.83-6.87 minute peak.

**Figure A3(a).**
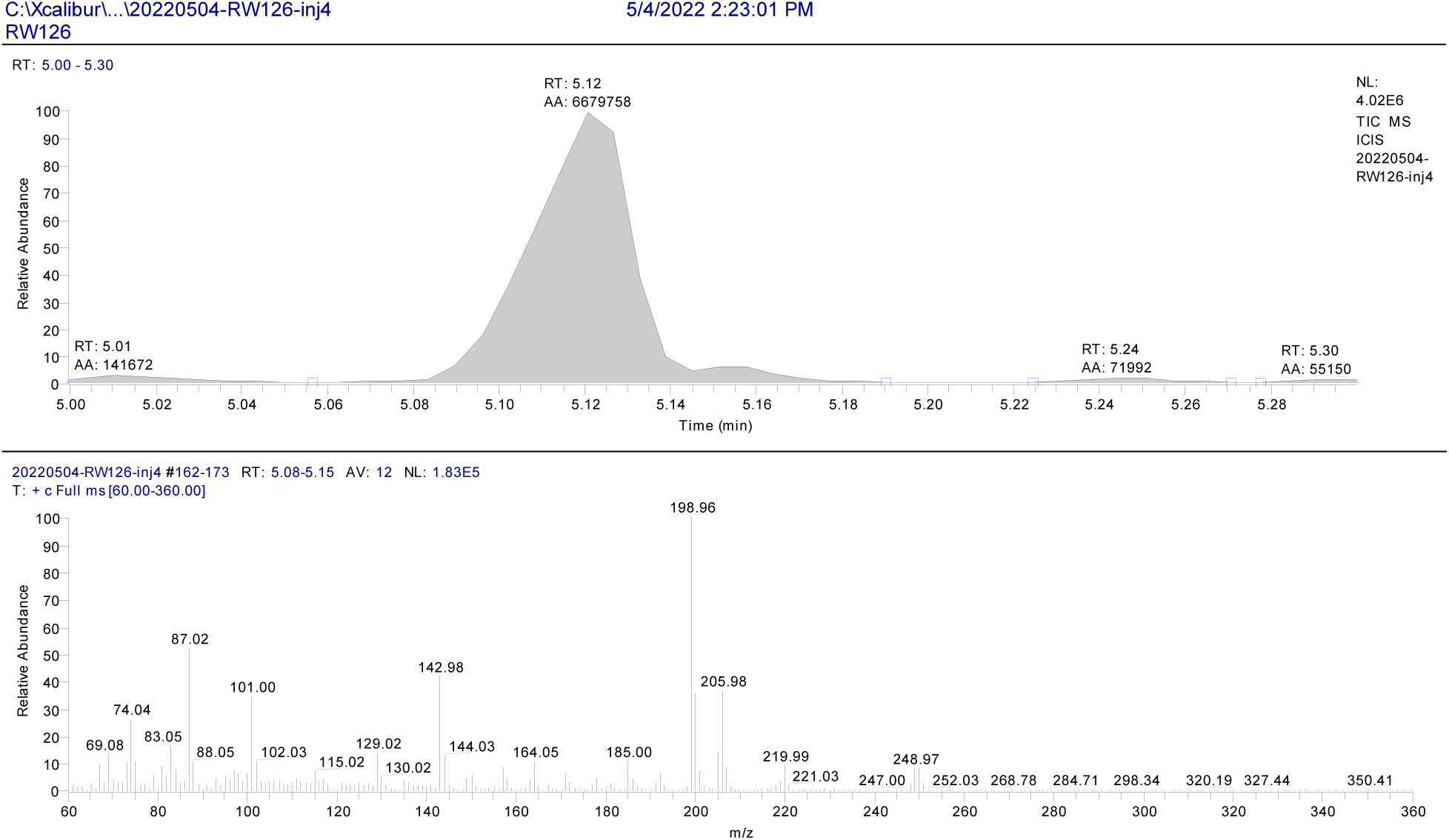
Chromatogram of the *X. campestris* iC14-d_7_ peak (“RT: 5.12”), and the iC14-d_7_ mass spectrum.

**Figure A3(b).**
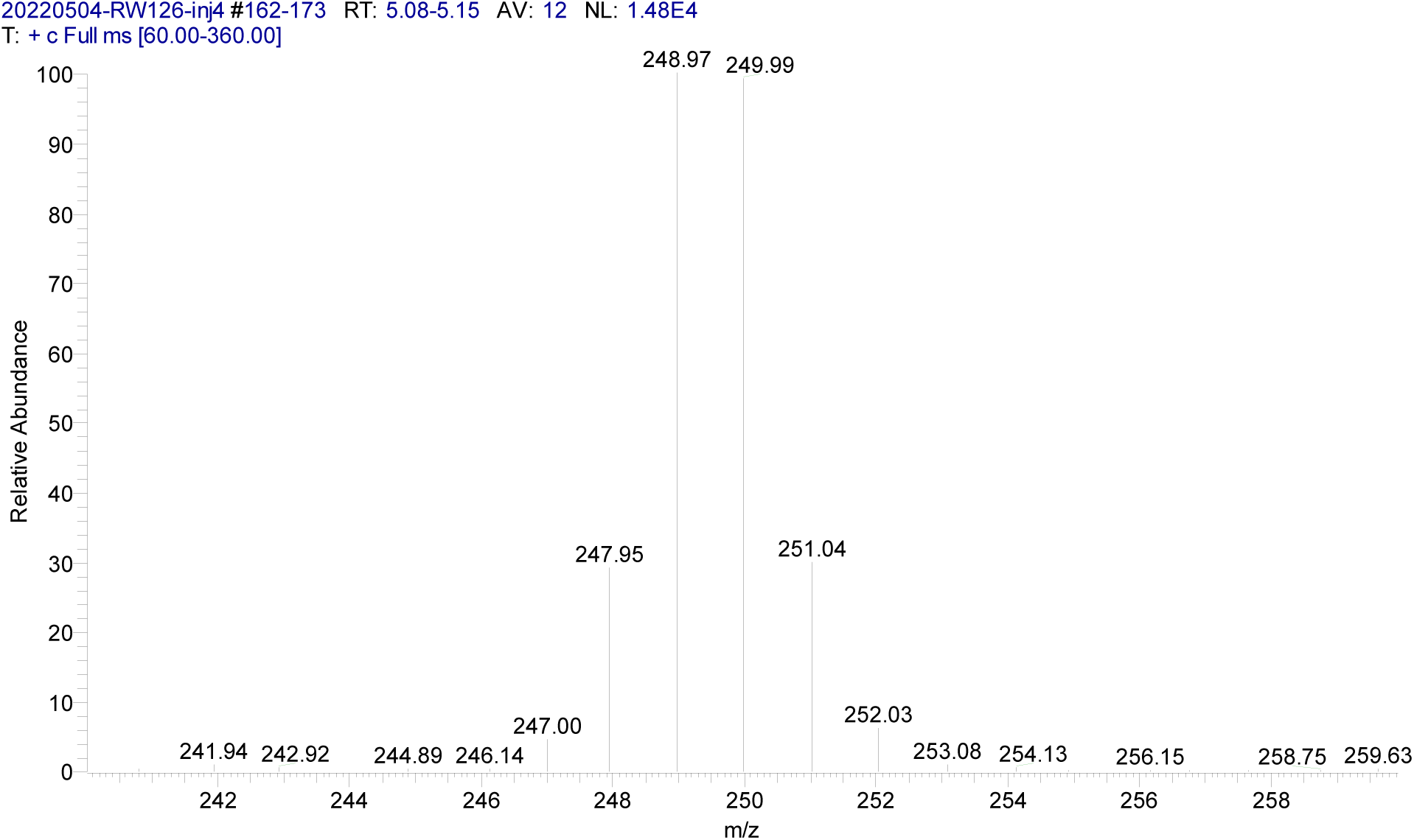
Detail of the mass spectrum of *X. campestris* iC14-d_7_.

**Figure A3(c).**
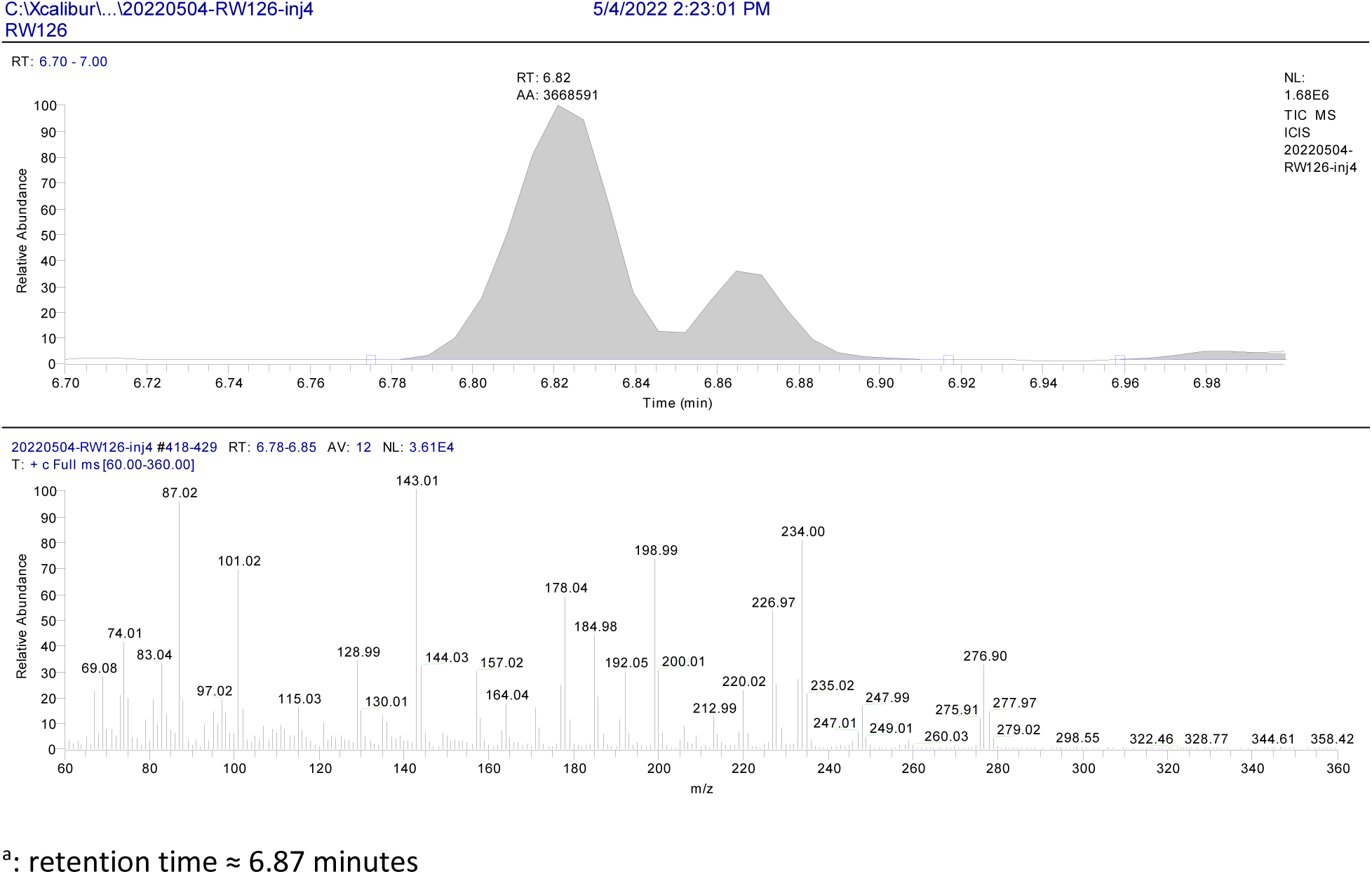
Chromatogram showing separation of *X. campestris* iC16-d_7_ (“RT: 6.82”) from iC16^a^, and the iC16-d_7_ mass spectrum.

**Figure A3(d).**
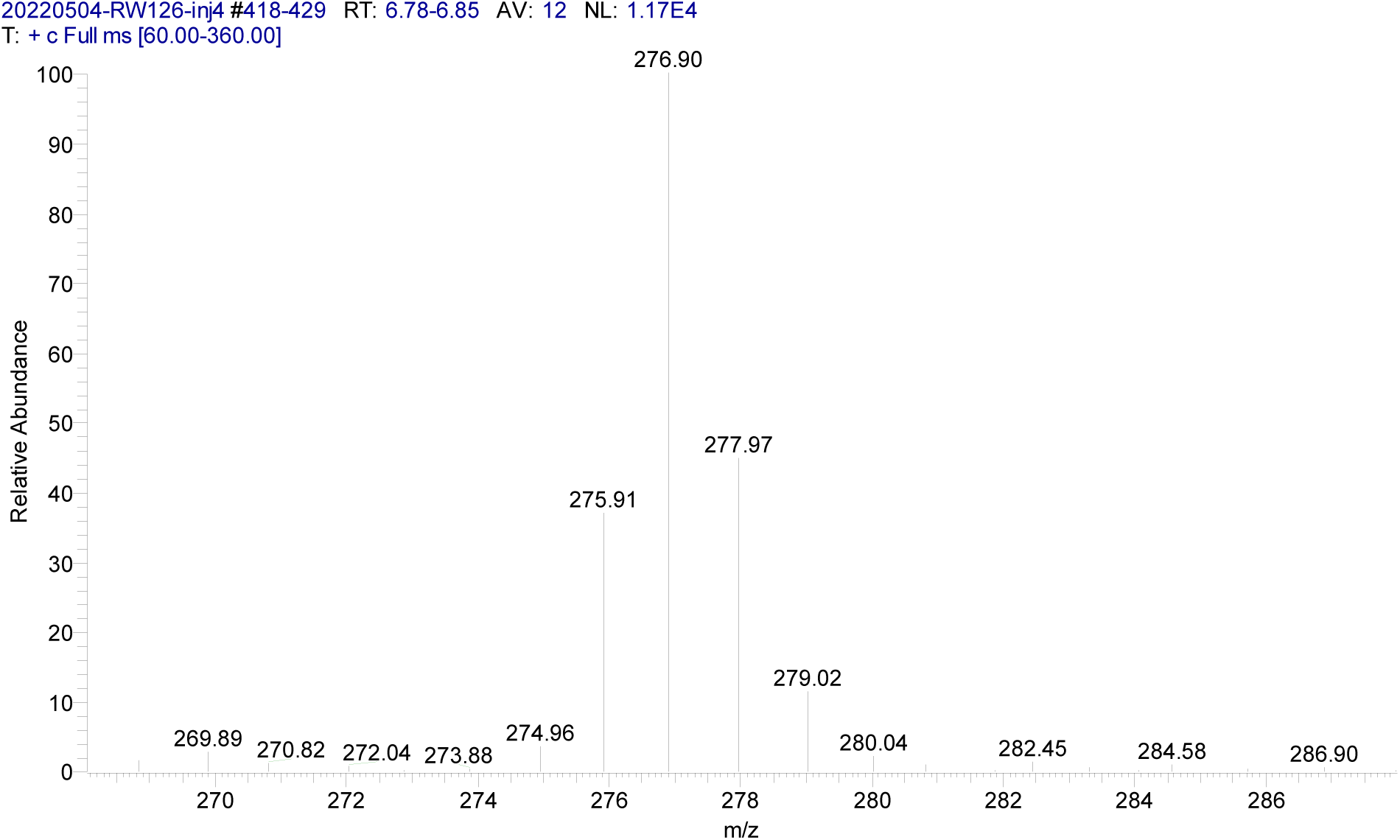
Detail of the mass spectrum of *X. campestris* iC16-d_7_.

**Figure A4.**
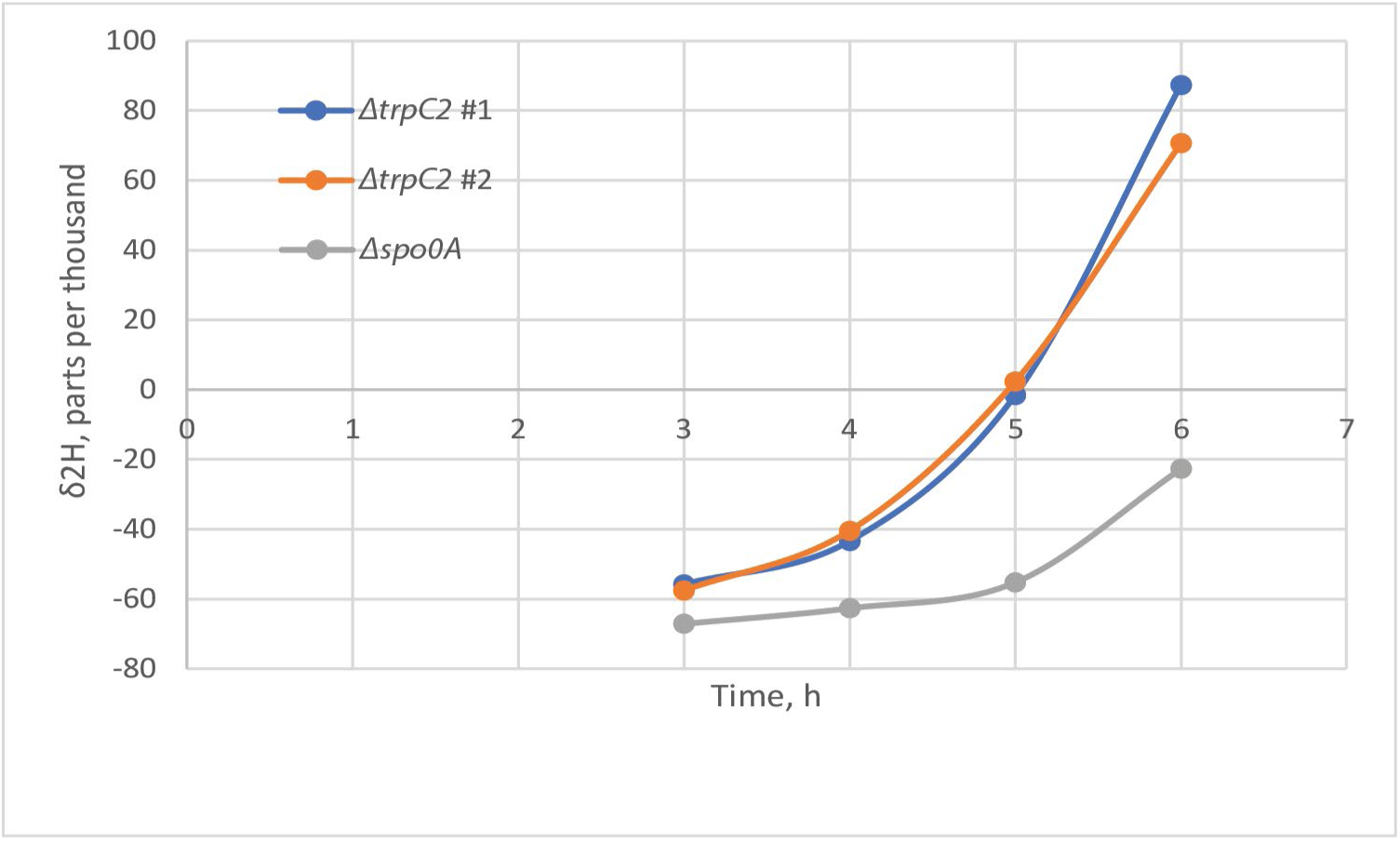
Broth water deuteration versus time in *B. subtilis ΔtrpC2* and *Δspo0A::ermtrpC2* cultures on sporulation medium supplemented with *L*-valine-d_8_. Each curve represents results for a single culture.

**Figure A5:**
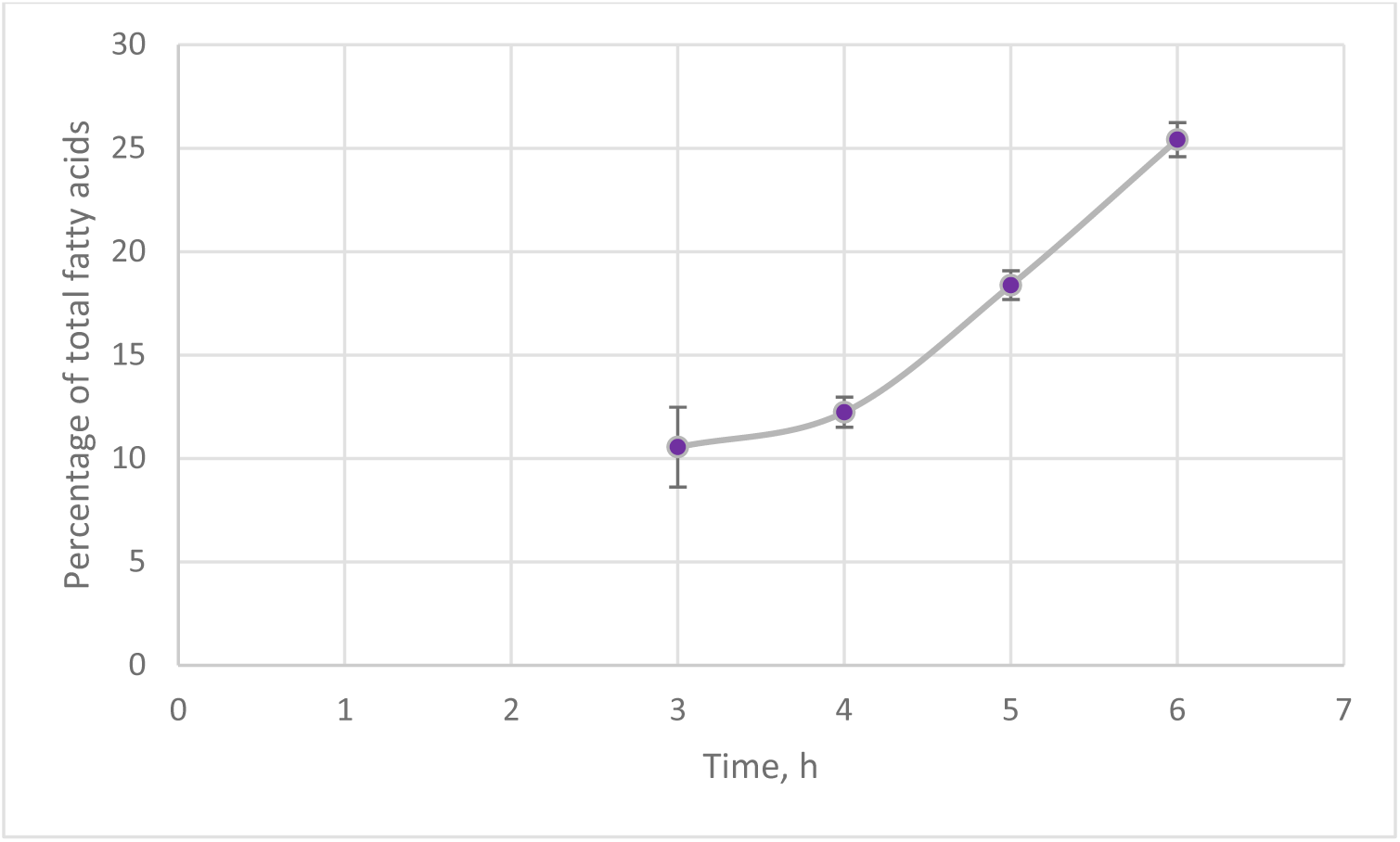
Percentage of iC15 fatty acid versus time for *B. subtilis Δspo0A::ermtrpC2* strain on sporulation medium supplemented with *L*-valine-d_8_. Error bars represent one SEM for duplicate cultures (n=2).

**Figure A6:**
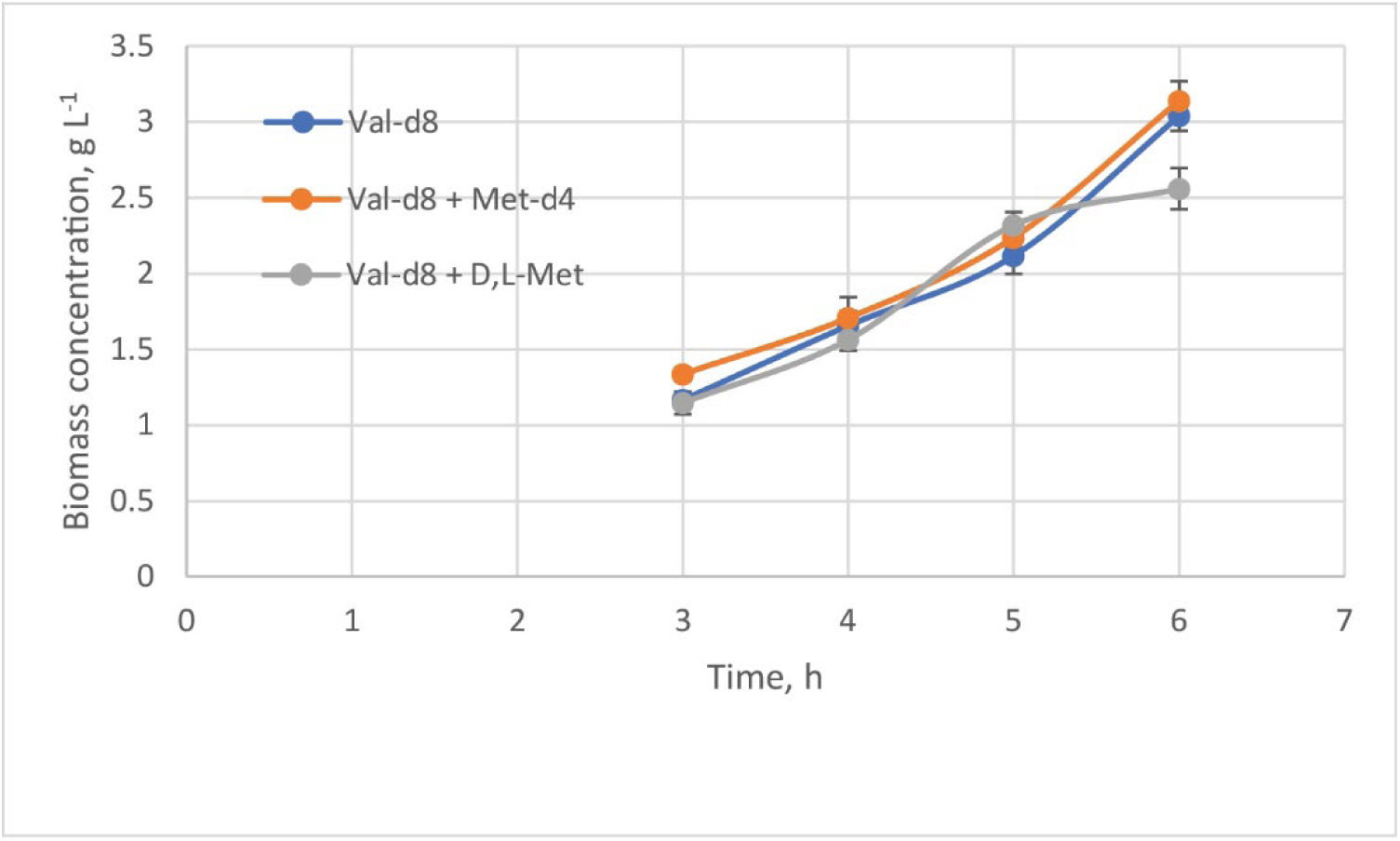
Biomass concentration versus time for triplicate *B. subtilis ΔtrpC2* sporulation medium cultures supplemented with *L*-valine-d_8_, *L*-valine-d_8_ and *D*,*L*-methionine-d_4_, or *L*-valine-d_8_ and *D*,*L*-methionine. Error bars represent one SEM for replicate cultures (n=3).

## Appendix 2

Summary of research on conversion of isobutyryl-CoA to propionyl-CoA by *B. subtilis*

Massey et al. (1976) described the usual five-step pathway for catabolism of isobutyryl-CoA to propionyl-CoA in bacteria. Based on genome-wide RNA profiling, Koburger et al. (2005) postulated one or more enzymes for each step, specifically those encoded by the *mmgABCDEF* and *fadNAE* operons, supplemented by *fadB* and *iolA*. The enzymes of the *mmgABCDEF* operon were first described by Bryan et al. (1996), and the function of the *fad* genes encoding fatty acid degradation enzymes was described by Matsuoka et al. (2007). Conversion by *B. subtilis* of methylmalonate semialdehyde to propionyl-CoA by IolA (formerly MmsA) was described by Stines-Chaumeil et al. (2006).

Koburger et al. (2005) postulated that FadE (formerly YusJ) and/or MmgC could dehydrogenate isobutyryl-CoA to methacrylyl-CoA, and Russell (2008) showed that *mmgC* encodes a short chain acyl-coenzyme A dehydrogenase (EC:1.3.99.-) with highest activity, amongst the substrates tested, against isobutyryl-CoA. The next step would presumably require the activity of an enoyl-CoA hydratase (EC:4.2.1.17) to form 3-hydroxyisobutyryl-CoA, which might be accomplished by FadB (formerly YsiB) or MmgB (Matsuoka et al., 2007). In the standard pathway, 3-hydroxyisobutyryl-CoA is hydrolyzed to the free acid then dehydrogenated to methylmalonic acid semialdehyde (Massey et al., 1976). How the latter would be formed is unclear, although MmgB possesses 3-hydroxybutanoyl-CoA dehydrogenase activity (EC:1.1.1.157; Vegunta, 2011); FadN (formerly YusL) is another possibility for at least part of this conversion (Matsuoka et al., 2007). The final step of decarboxylating and dehydrogenating the semialdehyde to propionyl-CoA is presumably accomplished by the methylmalonate semialdehyde dehydrogenase, IolA (EC:1.2.1.27; Stines-Chaumeil et al., 2006).

Our work with *ΔmmgA::ermtrpC2* and *ΔmmgC::ermtrpC2* strains grown on non-sporulation medium supplemented with *L*-valine-d_8_ showed that propionyl-d_5_-CoA (and propionyl-CoA) concentration was only partially reduced, indicating that MmgA and MmgC participate in isobutyryl-CoA catabolism but are replaceable by other enzymes, consistent with the report of Bryan et al. (1996) that deletion of all the genes from *mmgA* to *mmgD* had no obvious effect on growth or sporulation.

## Appendix 3

Proposed pathway from propionyl-CoA into iso-even fatty acids

Figure A7 shows a proposed pathway from propionyl-CoA to pimeloyl-ACP (-acyl carrier protein), which is assumed able to be integrated into the backbone of iso-even fatty acids, as described in detail below. An interesting aspect of the core segment of the pathway is its similarity to that used by *E. coli* to convert malonyl-CoA into pimeloyl-ACP as an intermediate of biotin synthesis (Lin et al., 2010). More importantly, *B. subtilis* has been shown to synthesize pimelic acid via the fatty acid synthesis pathway (Mandandhar and Cronan, 2017). In the following, it is assumed that the interconversion of any given acyl-CoA ester and the corresponding ACP ester (and possibly the free acid) occurs without difficulty.

We propose that propionyl-CoA is oxidized to malonyl-CoA via a four-stage process starting with its desaturation by acyl dehydrogenase, AcdA (UniProt P45867, EC:1.3.99.-), which is strongly expressed from t_2_ to t_5_ of sporulation (Nicolas et al., 2012; per SubtiWiki, www.subtiwiki.uni-goettingen.de, accessed March 22, 2023). The same reaction might also be achieved by FadE (UniProt O32176) of the fatty acid β-oxidation cycle (Fujita et al., 2007). The resulting acryloyl-CoA (or-ACP) would be hydrolyzed by enoyl-CoA hydratase FadB (UniProt P94549, EC:4.2.1.17) to form 3-hydroxy-propionyl-CoA, as occurs pathologically in humans (Ando et al., 1972). This alcohol would be oxidized to malonyl-CoA via the corresponding aldehyde through the action of alcohol dehydrogenase AdhB (UniProt O06012; Parro et al., 1997; Huyen et al., 2009), strongly expressed from t_2_ to t_5_ (Subtiwiki), and aldehyde dehydrogenase AdhA (UniProt C0SPA5, EC:1.1.1.-; *ibid.*).

The malonyl-ACP thus formed is proposed to be coupled to a second propionyl-CoA molecule by FabHA (UniProt O34746, EC:2.3.1.300), the β-ketoacyl-ACP synthase III initiating fatty acid biosynthesis in non-stressed *B. subtilis* cells. This enzyme has been shown to have activity with propionyl-CA similar to that with acetyl-CoA or isovaleryl-CoA as primer (Choi et al., 2000). The next three reactions comprise the well-known fatty acid synthesis cycle (Fujita et al., 2007) catalyzed by FabG (UniProt P51831, EC:1.1.1.100), YwpB (UniProt P94584, FabZ, EC:4.2.1.59) and FabI (UniProt P54616, EC:1.3.1.9). This would be followed by appendage of another malonyl-ACP unit to the growing fatty acid chain, catalyzed by β-ketoacyl-ACP synthase II, FabF (UniProt O34340, EC:2.3.1.179), and another synthesis cycle catalyzed by FabG, YwpB and FabI to form heptanoyl-ACP.

We propose that the heptanoyl-ACP formed in the above process is omega-oxidized by cytochrome P450/NADPH-P450 reductase, YetO (UniProt O08394, CypD, EC:1.14.14.1), purported to convert alkanes to primary alcohols, and most strongly expressed from t_2_ to t_5_ (SubtiWiki), although it has been shown to catalyze only sub-terminal oxidation of longer-chain fatty acids (Budde et al., 2005; Gustafsson et al., 2004). The primary alcohol hence formed would be sequentially oxidized to pimeloyl-ACP via the corresponding aldehyde by the reactions catalyzed by AdhB and AdhA described above.

Finally, although not shown in Figure A7, we propose that FabF is able to append a six-carbon segment of the pimeloyl-ACP thus formed into the iso-even fatty acids made with isobutyryl-CoA as primer.

**Figure A7.**
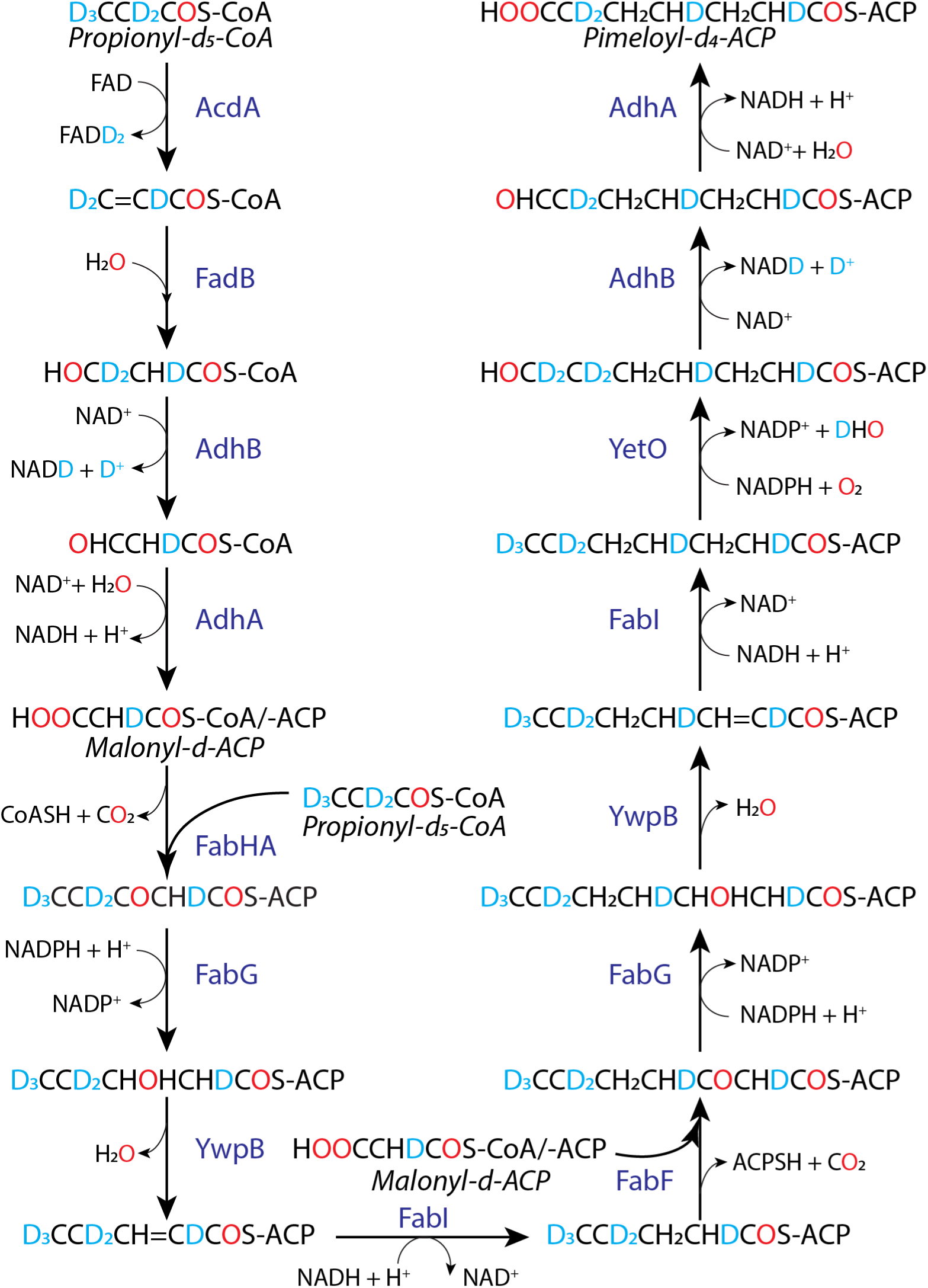
Proposed pathway from propionyl-d_5_-CoA to pimeloyl-d_4_-ACP. Transesterification reactions of acyl-CoA and ACP thioesters are not shown, and the details of deuterium and protium release are schematic. For cultures labeled with *L*-valine-^13^C_5_,^15^N, all carbon atoms explicitly shown are carbon-13.

